# Early life stress-induced miR-708-5p regulates mood disorder-associated behavioural phenotypes in mice and is a potential diagnostic biomarker for bipolar disorder

**DOI:** 10.1101/2024.03.14.584977

**Authors:** Carlotta Gilardi, Helena C. Martins, Alessandra Lo Bianco, Silvia Bicker, Pierre-Luc Germain, Fridolin Gross, Ayse Özge Sungur, Theresa M. Kisko, Frederike Stein, Susanne Meinert, Rainer K. W. Schwarting, Markus Wöhr, Udo Dannlowski, Tilo Kircher, Gerhard Schratt

## Abstract

Mood-disorders (MDs) are caused by a complex interplay of genetic and environmental (GxE) risk factors. However, the molecular pathways engaged by GxE risk factors to trigger specific MD-associated endophenotypes are still poorly understood. Here, by using unbiased small RNA sequencing in peripheral blood mononuclear cells (PBMCs), we identified the BD-associated miR-708-5p as one of the most strongly upregulated microRNAs in peripheral blood of both healthy human subjects with a high genetic or environmental (early life stress) predisposition to develop MDs. miR-708 is also upregulated in the hippocampus of rats which underwent juvenile social isolation, a rodent model of early life stress. Furthermore, ectopic overexpression of miR-708-5p in the hippocampus of adult male mice is sufficient to elicit MD-associated behavioural endophenotypes, demonstrating a causal role for elevated miR-708-5p levels in MD development. We further show that miR-708-5p directly targets Neuronatin (Nnat), an endoplasmic reticulum (ER) resident protein involved in calcium homeostasis. Consequently, restoring Nnat expression in the hippocampus of miR-708-5p overexpressing mice rescues miR-708-5p dependent behavioural phenotypes. Finally, miR-708-5p is strongly upregulated in PBMCs derived from patients diagnosed with MD, in particular BD males. Peripheral expression of miR-708-5p, in conjunction with the previously identified miR-499-5p, allows to differentiate male BD patients from patients suffering from major depressive disorder (MDD) and healthy controls. In summary, we describe a functional role for the miR-708-5p/Nnat pathway in MD etiology and identify miR-708-5p as a potential biomarker for the differential diagnosis of MDs.

## Introduction

MDs are a group of chronic psychiatric diseases affecting mood and cognition that include major depressive disorder (MDD) and bipolar disorder (BD). MDD is characterized by long, persistent, depressive episodes. BD has a distinct pattern of mood oscillations, from depressive episodes like MDD, and (hypo)manic phases. Furthermore, BD is subclassified in BD type 1 (mania and depression), and BD type 2 (hypomania and depression). It is known that MDs are highly heritable and share common genetic signature (McGuffin *et al*, 2003). In particular, BD is characterized by up to 70% heritability in monozygotic twins (Craddock & Sklar, 2013). However, complex polygenetic mechanisms cannot fully explain the onset of MD, and environmental factors, such as early life stress (ELS; e.g., physical and sexual abuse, emotional neglect), play an important role in the etiology and outcome of MD (Aas *et al*, 2020; Nemeroff, 2016; Rodriguez *et al*, 2021). Moreover, MD have a strong sex-specific component (e.g., higher prevalence of MDD, but not BD in females) (Noble, 2005) whose underlying biological mechanisms are unknown.

Although recapitulating the full spectrum of MD-associated symptoms is challenging, specific endophenotypes can be reliably assessed by employing dedicated behavioural tests in rodents. These include for example behavioural despair (FST, TST), anhedonia (sucrose preference), anxiety (OFT, EPM), cognitive impairments (mazes, NOR) and compulsive/manic-like behavior (marble burying) (Hoffman, 2013). Various chronic stress models have been established to mimic the impact of stress on MD symptomatology. Among them, post-weaning juvenile social isolation in rats represents a robust model to induce MD-associated endophenotypes, e.g., deficits in social communication and cognitive abilities (Seffer *et al*, 2015; Valluy *et al*, 2015). On the other hand, haploinsufficiency for the Cacna1c cross-disorder psychiatric risk gene is frequently used to study gene x environment interactions relevant for MD (Dedic *et al*, 2018). For example, *Cacna1c* (+/-) mice show decreased immobility in the tail suspension test (TST) and FST, higher preference for sucrose as well as decreased anxiety behavior, although the latter was a characteristic of female mice only (Dao *et al*, 2010). In Cacna1c (+/-) rats, a long-term environmental impact on object recognition, spatial memory and reversal learning was observed (Braun *et al*, 2019).

microRNAs (miRNAs) are a large family of small (∼22nt), noncoding RNAs that act as posttranscriptional regulators by binding to complementary sequences in the 3’-untranslated region (UTR) of target messenger RNAs (mRNAs) (15) (Bartel, 2018). Recent research has highlighted the role of miRNAs in the pathogenesis of MDs (Martins & Schratt, 2021). Main pathways affected by miRNA dysregulation in the context of MD animal models include serotonergic neurotransmission (e.g., miR-16, miR-34), glucocorticoid signaling (e.g., miR-17-92 cluster, miR-15), neurotrophins (e.g., miR-182), Wnt signaling (e.g., miR-124) and synaptic plasticity (e.g., miR-134, miR-218). While these candidate studies in rodents provided important insight into biological mechanisms, the translational value for MD therapeutics and diagnostics is mostly limited by the lack of corresponding human data.

In humans, miRNAs have been associated with MD etiology in expression studies of postmortem brain tissue (Moreau *et al*, 2011) and blood samples from living patients (Dwivedi, 2011). Differential expression of miRNAs in peripheral blood mononuclear cells (PBMCs) has been previously investigated for biomarker discovery in MD. For example, miR-499-5p was significantly upregulated in female and male PBMC samples of BD (Martins *et al*, 2022). miR-124-3p was found to be significantly upregulated in PBMCs samples of MDD compared to healthy controls and it was downregulated after eight weeks of antidepressant treatment (He *et al*, 2016). However, the overlap between these studies is usually low, and therefore these attempts so far have not yielded reliable miRNA biomarkers in MDs.

We therefore decided to undertake an unbiased, back-translational approach to identify MD-associated miRNAs from a large human cohort of healthy subjects at high genetic and environmental risk to develop MD. One of the identified GxE regulated miRNAs, the BD-associated miR-708-5p (Forstner *et al*, 2015), was further functionally characterized in rodent models and subsequently tested for its diagnostic potential in MD patient subgroups.

## Materials and Methods

### Human study

#### Recruitment of Participants

The study involved participants with BD (26 females, 37 males), MDD (18 females, 24 males), and healthy individuals (26 females, 31 males), including psychiatrically healthy subjects that had a history of childhood maltreatment (17) or a genetic predisposition to MDs (18) or no risk (18). These participants were recruited from the University of Marburg and the University of Münster in Germany, as part of the FOR2107 cohort (39) (Kircher *et al*, 2019). Diagnoses were made using the SCID-I interview (Wittchen, 1997), and adapted to DSM-IV criteria, excluding those with substance abuse, severe neurological, or other significant medical conditions. Healthy controls were screened similarly, with inclusion in the maltreatment study if at least one of the subclasses of the Childhood Trauma Questionnaire (CTQ) reached the maltreatment threshold (Emotional Abuse ≥ 10, Physical Abuse ≥ 8, Sexual Abuse ≥ 8, Emotional Neglect ≥ 15, and Physical Neglect ≥ 8) (Walker *et al*, 1999). All participants underwent a comprehensive neuropsychological test battery, including the d2 test of attention (Brickenkamp, 2002). All necessary ethical approvals were obtained (ethics committees of the Medical Faculties of the Universities of Münster (2014-422-b-S) and Marburg (AZ: 07/14)), and participants consented to the study, which complied with the Declaration of Helsinki and the Belmont Report. Demographic and clinical data are outlined in Table 1 and in Supplementary Table 1.

**Table 1.**

|  | **Control** | **BD** | **MDD** | **P Value** |
| --- | --- | --- | --- | --- |
| *n (f/m)* | 26/31 | 26/37 | 18/24 | N/A |
| Age ± S.D. | 29.4 ± 5.4/<br>39.7 ± 13.4 | 29.9 ± 5.4/<br>41.7 ± 11.3 | 29.6 ± 4.9/<br>39.3.6 ± 14.7 | 0.94/0.75 |
| CTQ ± S.D. (%) | 40.7 ± 11.4<br>(53.8%) | 42.3 ± 13.2<br>(30.8%) | 50.2 ± 20.2<br>(61%) | 0.1067/0.4170 |
| Family history<br>of MDs (%) | 8 (30.8%)/<br>5 (16.1%) | 10 (38.5%)/<br>10 (27.0%) | 5 (27.8%)/<br>7 (29.1%) | N/A |
| HAMD ± S.D. | 2.7 ± 3.9/<br>1.5 ± 2.5 | 7.5 ± 6.2/<br>8.6 ± 6.4 | 14.8 ± 6.7/<br>10.3 ± 7.7 | <0.0001 |
| YMRS ± S.D. | 0.5 ± 1.1/<br>0.7 ± 1.7 | 2.4 ± 2.9/<br>7.0 ± 8.2 | 1.4 ± 1.6/<br>1.7 ± 2.2 | <0.01/<br><0.0001 |
| BDI ± S.D. | 6.1 ± 6.1/<br>4.5 ± 3.9 | 12.9 ± 11.1/<br>12.9 ± 9.6 | 26.6 ± 11.0/<br>16.6 ± 10.7 | <0.0001 |
| AD (%) | 0 (0%) | 7 (26.9%)/<br>17 (45.95%) | 16 (88.9%)/<br>17 (70.83%) | N/A |
| Antipsychotic<br>(%) | 0 (0%) | 12 (46.15%)/<br>19 (51.35%) | 7 (38.89%)/<br>4 (16.67%) | N/A |
| Lithium (%) | 0 (0%) | 4 (15.38%)/<br>12 (32.43%) | 1 (5.56%)/<br>0 (0%) | N/A |
| Anticonvulsive<br>(%) | 0 (0%) | 6 (23.08%)/<br>10 (27.03%) | 1 (5.56%)/<br>1 (4.17%) | N/A |
| Stimulants (%) | 0 (0%) | 0 (0%)/<br>1 (2.70%) | 0 (0%)/<br>2 (8.33%) | N/A |
| Benzodiazepin<br>e (%) | 0 (0%) | 0 (0%) | 1 (5.56%)/<br>0 (0%) | N/A |
| Z substance<br>(%) | 0 (0%) | 0 (0%)/<br>1 (2.70%) | 1 (5.56%) | N/A |

Subjects for miR-708-5p expression analysis on psychiatrically healthy controls (Control), Bipolar disorder patients (BD) or Major Depressive Disorder patients (MDD). One-way-ANOVA was performed to evaluate significant differences between groups. S.D.: standard deviation, CTQ: Childhood Maltreatment

#### Human peripheral blood mononuclear cell (PBMC) sample processing

PBMCs were obtained from 10 ml of whole blood using the LeukoLOCK technology (Thermo Scientific) at the Biomaterialbank Marburg, Germany. After sample randomization, total RNA extraction from PBMCs was carried out using the *mir*Vana™ kit (Thermo Fisher) or TRIzol™ Reagent (Thermo Fisher) following the manufacturer’s protocol.

#### PBMCs processing for small RNA-sequencing

For the Small RNA-sequencing, RNA was extracted with *mir*Vana™ kit (Thermo Fisher) and DNase treated, and it was carried out at Functional Genomic Center Zurich (FGCZ, https://fgcz.ch/ ).

#### Small RNA-analysis

Short RNA reads were processed using the ncPro 1.6.4 pipeline (Chen *et al*, 2012) using the hg19 annotated (based on miRBase version 21). Only mature miRNA reads (accepting +2bp on either end) were considered for downstream analysis. Differential expression analysis was then performed using limma/voom 3.46.0 (Ritchie *et al*, 2015), with a model including a covariate to correct for the two technical batches.

### Rat primary neuron culture

Cultures of primary hippocampal and cortical neurons from rat hippocampus were established using E18 Sprague-Dawley rats obtained from Janvier Laboratories, as described previously (Martins *et al*., 2022).

#### Transfection

Primary hippocampal neurons were transfected with Lipofectamine™ 2000 Reagent and plasmid DNA constructs as previously described (Martins & Schratt, 2021).

#### Virus infection

Primary hippocampal neurons were infected with recombinant adeno-associated virus (rAAV) containing hpCTL or hp708 constructs on *days in vitro* (DIV) 2 by adding 300μl of a mixture of the virus with 300µl of NBP+ per well of a 24-well plate. Cells were used for downstream analysis on DIV19-20.

#### Luciferase Assay

DIV6 hippocampal neurons were transfected with 100ng of the Nnat-3’UTR luciferase reporters and 500ng of hpCTL-p or hp708-p, spCTL-p or sp708-p. After seven days from transfection, the cells were lysed, luciferase assay was conducted using a GloMax R96 Microplate Luminometer (Promega), as previously described (Martins *et al*., 2022). The relative luciferase activity was determined by calculating the ratio of the Firefly signal to the Renilla signal.

### *In Situ* Hybridization

Single-molecule fluorescence *in situ* hybridization (smFISH) for miRNA detection in hippocampal neuron cultures was conducted using the QuantiGene ViewRNA miRNA Cell Assay Kit (Thermo Fisher, QVCM0001) following the manufacturer protocol with minor adjustments. Probes for has-miR-708-5p (Alexa Fluor 546, Thermo Fisher), Nnat (Alexa Fluor 546), CamK2 (Alexa Fluor 488) and Gad2 (Alexa Fluor 488) were used for the assay.

#### Plasmid design and rAAVs preparation

For rAAV-mediated overexpression of miR-708-5p, the chimeric miR-708 hairpin was generated by polynucleotide cloning. A detailed description and characterization of this system have been published (Christensen *et al*, 2010). To knockdown miR-708-5p, a sponge plasmid was constructed by inserting six TDMD sites predicted to bind miR-708-5p in the 3’UTR of eGFP on rAAV-hSyn-EGFP. To perform luciferase activity analysis, the wild type and mutated Nnat 3’UTR was inserted into a pmirGLO dual-luciferase expression vector (Promega). Detailed description of the design of these constructs and of the rescue constructs are listed in the supplementary material section. Viral vectors were produced by the Viral Vector Facility (VVF) of the Neuroscience Center Zurich (https://www.vvf.uzh.ch).

### Animal studies

#### Mouse experiments

##### Husbandry, housing, and behavioral testing

All animal experiments on mice were conducted in accordance with Switzerland’s animal protection laws and received approval from local cantonal authorities (Approval ID: ZH194/21). Mice were housed collectively in cages designed for 2 to 4 individuals, with unrestricted access to food and water. The animal facility maintained an inverted light-dark cycle of 12 hours, with behavioral assessments conducted during the dark phase. Prior to experimentation, mice underwent daily handling lasting 5 minutes each over one week. Details about general behavioral procedures can be found in the supplementary materials and methods section.

##### Stereotactic surgeries and post-operative care

Stereotactic brain injections were conducted on 2-month-old C57BL/6JRj wild-type mice. The mice were anesthetized with 5% isoflurane in oxygen (1 L/min) and positioned on a stereotactic frame. Viral injections were carried out bilaterally. For dorsal hippocampus, the coordinates used were AP: -2.1mm; ML: ±1.5mm. For ventral hippocampus, the coordinates used were AP: -3.3mm; ML: ±2.7mm. Detailed description of the procedure and post-operative care can be found in the supplementary materials and methods section.

##### Open Field Test

Mice were placed in an open field box (size: L 45 x W 45 x H 40 cm, TSE System, Bad Homburg, Germany) with a 60-minute exploration window. Dim, yellow light was used in the Open field test of Fig. S2H and I, and Fig. S4C, D, E. 30 lux, white light was used for the Open Field in Fig. S2J. The experiment was recorded, and the TSE VideoMot2 analyzer software (TSE Systems, Bad Homburg, Germany) was used for the automated assessment.

##### Elevated Plus Maze Test

The mouse was placed on the center of the elevated plus maze, a cross-shaped apparatus with two open arms and two closed arms, each arm measuring L 65 x W 5.5 cm and elevated 62 cm from the ground. 20 lux white light was used. Behavior was recorded on video for five minutes. The videos were automatically analyzed using TSE VideoMot2 analyzer software (TSE Systems, Bad Homburg, Germany)

##### Saccharin Preference Test

Mice were habituated in their standard group housing to the presence of two bottles for 48 hours before the test. The test was performed by single housing the test animals, and two bottles of tap water or 0.1% Saccharin solution were given to them for 48 hours. The weight of the bottles was recorded at the beginning of the test and every 12 hours around the light change (09:00-09:15 am/pm). The position of the bottle was exchanged each time to avoid side preference. The animals were re-grouped after the test ended.

##### Novel Object Recognition (NOR) Test

The test was performed as previously described (Daswani *et al*, 2022), with slight modifications: 1. Twenty-four hours prior to testing, the animals were habituated to the arena used for the test; 2. The break between familiarization and novelty introduction rounds was extended to five minutes or 24 hours.

##### Marble Burying Test

The test was conducted as previously described (Levone *et al*, 2021).

##### Passive Avoidance Test

The test was conducted using the Passive Avoidance 2-Compartment light-dark arena of the TSE System. The white light was set to 500 lux. On day one, the animal was placed into the light compartment while the door to the dark one was open and it was allowed to explore; once it passed to the dark compartment, the door closed, and the animal received a 2-seconds foot shock of 0.3 mA after three seconds. On day two, the animal was placed again into the light compartment and, after 15 seconds, the door was automatically opened. The latency to enter the dark compartment was measured.

##### Tail Suspension Test

The mouse was hung by its tail for a duration of six minutes using a piece of tape affixed to the suspension metal bar, elevated approximately 50 cm from the ground. Before initiating the experiment, a climb-stopper, following a published procedure (Can *et al*, 2012), was placed at the mouse tail base to prevent climbing. The sessions were recorded on video, and the duration of immobility was subsequently quantified.

##### Tissue collection

Animals were sacrificed by cervical dislocation and the hippocampus was collected on an ice-cold glass plate and subsequentially snap-frozen for RNA extraction (Trizol protocol) and gene expression analysis or for histological assessments after 4% paraformaldehyde fixation, cryoprotection through 30% sucrose solution, and coronal sectioning at the cryostat. Further details can be found in the supplementary materials and methods section.

### Rat experiments

All animal experiments on rats were conducted in accordance with the National Institutes of Health Guidelines for the Care and Use of Laboratory Animals and were subject to prior authorization by the local government (MR 20/35 Nr. 19/2014 and G48/2019; Tierschutzbehörde, Regierungspräsdium Giessen, Germany). Constitutive heterozygous *Cacna1c^+/−^* animal breeding and the juvenile social isolation paradigm were conducted as described (Martins *et al*., 2022).

### RT-qPCR

Total RNA extraction from mouse brain tissue, PBMC samples, and primary hippocampal cultures was performed using TRIzol™ Reagent (Thermo Fisher, 15596026), and total RNA was subsequently extracted following the manufacturer’s guidelines. Further details can be found in the supplementary materials and methods section.

### PolyA RNA sequencing sample preparation

Total RNA extraction from mouse hippocampus was extracted TRIzol™ Reagent (Thermo Fisher, 15596026). The RNA was then treated with Turbo DNase enzyme. 600 ng of DNase-treated RNA was used for High Throughput Transcriptome sequencing. Libraries were prepared with Illumina TrueSeq mRNA protocol and the transcriptome sequencing was run on Illumina Novaseq 6000. The transcriptome sequencing was carried out by the Functional Genomic Center Zurich (FGCZ, hyperlink: https://fgcz.ch/ ).

### PolyA RNA sequencing analysis

Reads were mapped to the GRCm39 genome with STAR 2.7.8a (Dobin *et al*, 2013) using the GENCODE M26 annotation as reference and quantified at the gene-level using featureCounts 1.6.4 (Liao *et al*, 2014). Genes were filtered using edgeR’s filterByExpr function before differential expression analysis with edgeR 3.32.1 (Robinson *et al*, 2010) using likelihood ratio tests with 3 surrogate variables estimated using sva 3.38.0 (Leek *et al*, 2012). TargetScan 7 (Agarwal *et al*, 2015) was used to predict miRNA targets. For gene expression across different brain cell types (Figure 4B), the Allen 10X+smartSeq taxonomy and per-cluster expression was used, aggregating non-neuronal cell types into broader classes.

### Western blot

Proteins from primary hippocampal neurons were isolated using ice cold RIPA lysis buffer. 20ug of protein was mixed with 4xLaemmli Sample Buffer (Biorad) and were run on a 4–20% Mini-PROTEAN® TGX™ Precast Protein Gels (Biorad). Proteins were transferred on a nitrocellulose membrane and blocked for two hours at room temperature in blocking solution and incubated in primary antibody solution for 48 hours at 4°C. Membranes were washed and incubated with secondary antibody for 1h. Membranes were washed five times in TBS-T, developed with the Clarity™ Western ECL Substrate (Bio-Rad) and visualized with the ChemiDocTM MP, Imaging System (BioRad). Further details can be found in the supplementary materials and methods section.

### Statistical analysis

Details regarding the statistical analysis are listed in the Supplementary Materials and methods section.

## Results

### miR-708-5p is upregulated in the peripheral blood of human healthy subjects harboring an elevated genetic and environmental risk for MD

We hypothesized that miRNAs whose expression correlates with environmental (ER) and genetic risk (GR) for MD in human subjects might represent strong candidates for miRNAs causally involved in MD etiology. Furthermore, by starting from a human cohort, we hoped to identify miRNA candidates with high translational potential, meaning that they could serve as targets for miRNA therapeutics and/or diagnostics in MD.

To identify candidate miRNAs in an unbiased manner, we performed small-RNA sequencing with total RNA obtained from peripheral blood mononuclear cells (PBMCs) of healthy subjects (n=52) from the FOR2107 cohort (39)(Kircher *et al*., 2019) characterized by a high genetic (at least one first degree relative diagnosed with a mood disorder; GR group; n=14) or environmental (ER group; childhood trauma based on childhood trauma questionnaire (CTQ) score; n=16) predisposition to develop MDs (Fig. 1A; Table 1). For this initial analysis, we selected females since we were able to form more homogeneous GR and ER groups with females compared to males. We then focused on miRNAs that were differentially expressed in both ER and GR compared to healthy control subjects (CTL group; no known genetic or environmental risk factors; n=18). We found that a total of six miRNAs (miR-412-5p, miR-100-5p, miR-501-3p, miR-642a-5p, miR-4999-5p, miR-708-5p) fulfilled this criterium (Fig. 1B and C). Among them miR-708-5p was of specific interest, since it was the most strongly upregulated miRNA. Furthermore, it was previously shown to be induced by various forms of cellular stress (26–30) (Behrman *et al*, 2011; Lin *et al*, 2015; McIlwraith *et al*, 2022; Rodriguez-Comas *et al*, 2017; Yang *et al*, 2015) and had been associated with BD in the past (21)(Forstner *et al*., 2015). We could confirm that miR-708-5p is significantly upregulated in the GR and ER groups using qPCR (Fig. 1D). Thus, we decided to focus on miR-708-5p for our further studies.

**Fig. 1.**
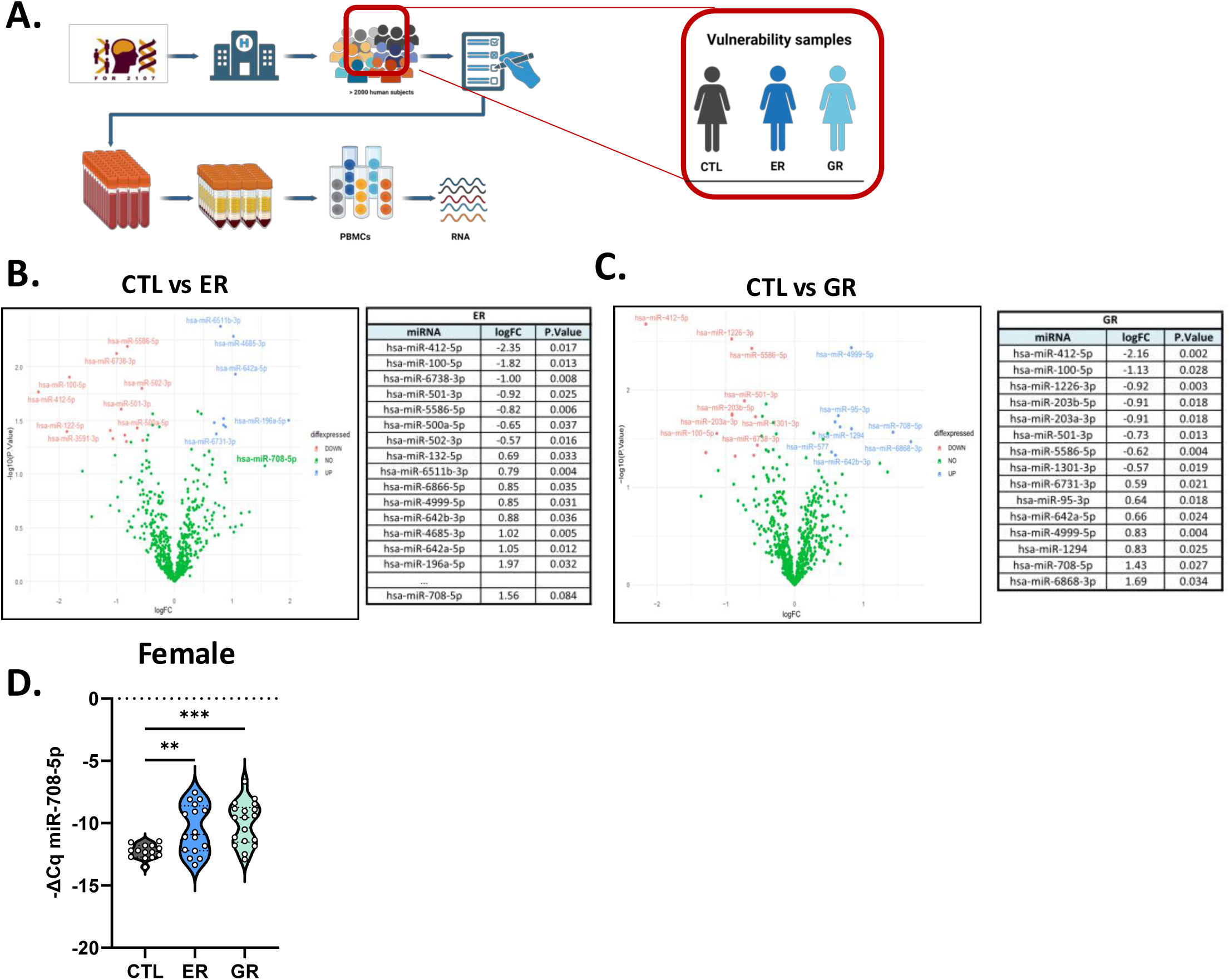
miR-708-5p is upregulated in the peripheral blood of human subjects at risk for with MDs. **A.** Schematic representation of the workflow to obtain PBMCs samples from the FOR2107 cohort (see also materials and methods) (39). Subjects underwent psychiatric and cognitive evaluation. After the data was collected, blood was withdrawn and PBMCs prepared followed by RNA extraction and downstream gene expression analysis. Samples were collected from psychiatrically healthy female subjects (CTL), or psychiatrically healthy female subjects with genetic predisposition for mood disorder (GR) or that suffered from childhood maltreatment (ER) (Vulnerability samples). **B. and C.** Volcano plot depicting miRNAs differentially expressed based on small RNA sequencing performed with RNA isolated from PBMCs of control subjects (CTL; n=18), healthy subjects who suffered childhood maltreatment (Environmental risk, ER; n=18), or healthy subjects with a genetic risk (GR; n=14). CTL vs ER (B) or GR (C): miRNAs that are significantly downregulated (logFC<0.5, p-value<0.05) are marked in red; miRNAs that are significantly upregulated (logFC>0.5, p-value<0.05) in blue. Each Volcano Plot is accompanied by a table showing the top 15 differentially expressed genes ordered from smallest to largest p-value. miRNAs overlapping between (B) and (C) are highlighted. **C.** miR-708-5p qPCR analysis of total RNA isolated from PBMCs of control subjects (CTL=18), or healthy subjects who suffered childhood Maltreatment (Environmental risk, ER= 16), or healthy subjects with a genetic risk (GR=18). One-way ANOVA with Holm-Sidak’s multiple comparisons test, HC vs ER, **, P=0.0064; HC vs GR, ***, P=0.0006; ER vs GR, ns. Data are presented as violin plots with median, quartiles and data points.

### miR-708-5p is expressed in rat hippocampal neurons and upregulated in the hippocampus of rat models of environmental or genetic risk for MDs

To further study the functional role of miR-708-5p in MDs, we considered rodents given their extensive use in modelling psychiatric conditions. As a first step, we investigated whether miR-708-5p was expressed in different regions of the rat brain (Fig. 2A). We observed robust expression in regions classically implicated in MDs and cognitive function, such as the amygdala, frontal cortex, and hippocampus (Fig. 2A). Next, we assessed miR-708-5p expression in rat hippocampal primary neurons. Single-molecule fluorescence *in situ* hybridization (sm-FISH) showed that miR-708-5p is expressed in both CamK2a+ excitatory and GAD2+ inhibitory neurons in rat hippocampal cultures, illustrating its widespread neuronal distribution (Fig. 2B and 2C). miR-708-5p expression in rat hippocampal cultures decreased over time during *in vitro* development, indicating a role for miR-708-5p at early stages of neuronal development (Fig. 2D). Motivated by the findings of miR-708-5p elevation in peripheral blood of humans at risk to develop MDs (Fig. 1B-D), we asked whether miR-708-5p was similarly dysregulated in rat genetic and environmental models of MDs. As a genetic model, we chose rats heterozygous for *Cacna1c* (*Cacna1c^+/-^),* a repeatedly validated cross-disorder psychiatric risk gene (60–65) (Bhat *et al*, 2012; Dao *et al*., 2010; Dedic *et al*., 2018; Harrison *et al*, 2022). Environmental risk was modelled by juvenile social isolation, a widely recognized model for early life trauma, e.g., childhood maltreatment (66) (Seffer *et al*., 2015) (Fig. 2E). miR-708-5p was significantly upregulated by social isolation in the hippocampus of juvenile wild-type rats compared to their group-housed counterparts (Fig. 2F), indicating parallels between the early life stress-related modulation of miR-708-5p expression in human peripheral blood and rat brain. Furthermore, *Cacna1c* heterozygosity led to an upregulation of miR-708-5p in the rat hippocampus independent of social housing conditions, suggesting that, like our human PBMC data (Fig. 1C-D), a *genetic* predisposition for MDs is sufficient to induce miR-708-5p expression in the rat brain (Fig. 2F). Our comprehensive analysis across various experimental paradigms revealed that miR-708-5p expression is linked to both environmental stressors and genetic factors associated with MDs in both humans and rodents. Therefore, manipulation of miR-708-5p in the rodent brain represents a viable strategy to study the involvement of miR-708-5p in the development of mood disorder-associated endophenotypes as well as the underlying molecular mechanisms.

**Figure 2.**
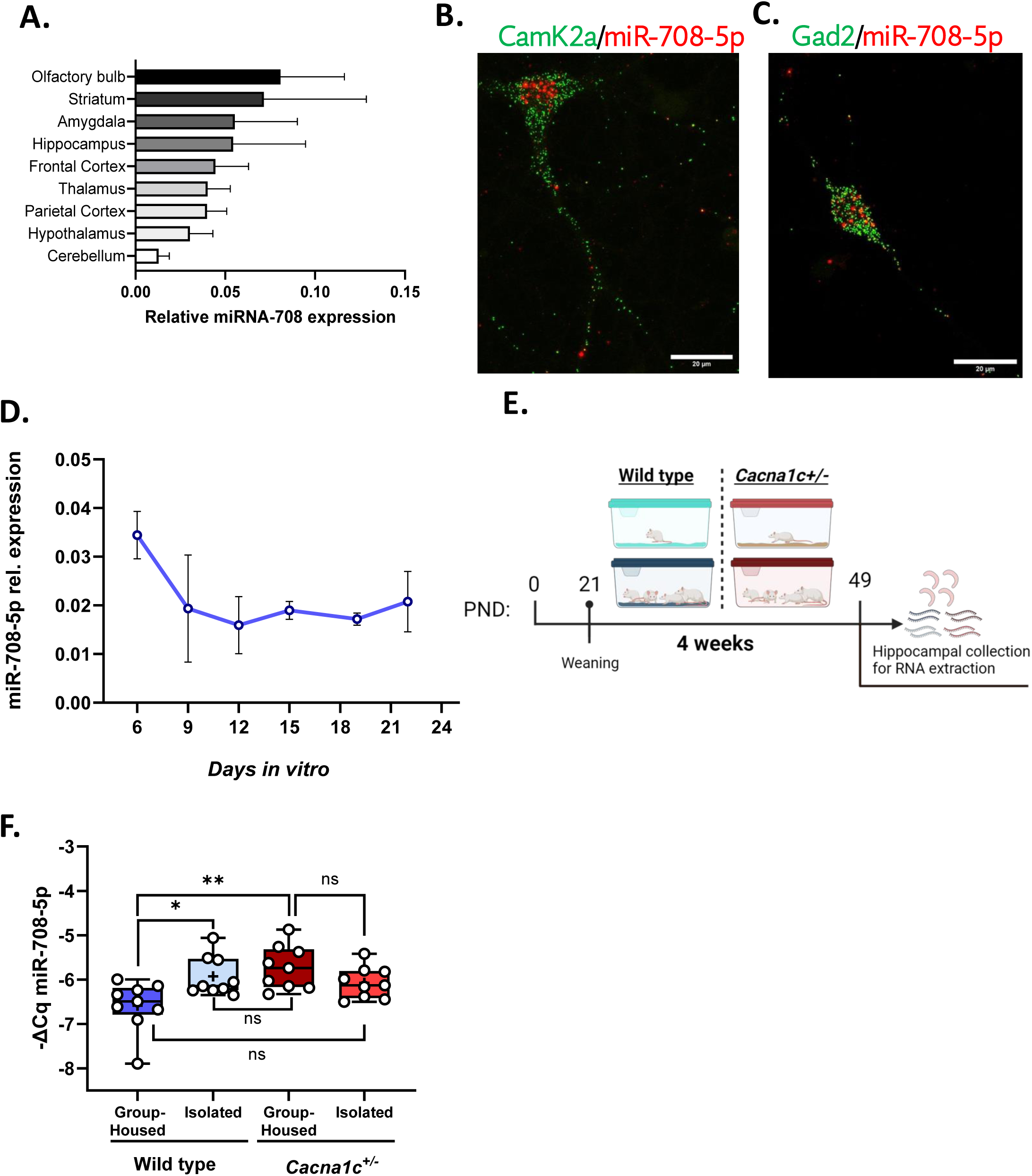
miR-708-5p is expressed in rat hippocampal neurons and upregulated in the hippocampus of rat models of environmental or genetic risk for MDs. **A.** miR-708-5p qPCR analysis using total RNA isolated from different areas of the adult female rat brain. Data are represented as bar graphs, mean ± SD (n=3 animals). **B.** Single-molecule fluorescence *in situ* hybridization (smFISH) performed in rat hippocampal neurons at 7 days *in vitro* (DIV7) using probes directed against miR-708-5p (red channel) and CamK2a (green channel) to identify excitatory neurons. Scale bar: 20μm. **C.** smFISH performed in rat hippocampal neurons at DIV7 using probes against miR-708-5p (red channel) and Gad2 (green channel) to identify inhibitory neurons. Scale bar: 20μm. **D.** Relative expression of miR-708-5p in primary rat hippocampal neurons at different DIV (n = 3 independent experiments) normalized to U6 snRNA. Data are represented on XY graph as mean ±SD. **E.** Schematic representation of the juvenile social isolation experiment, n=9 for each experimental group; PND: postnatal day. **F.** miR-708-5p qPCR analysis of total RNA isolated from hippocampi of Wild type (WT) or *Cacna1c^+/-^* male juvenile rats that were either group-housed or socially isolated for four weeks (n=9 in each group). WT group-housed vs WT Isolated: *, p=0.0265; WT group-housed vs *Cacna1c*^+/-^: **, p=0.0026; linear mixed model, - DeltaCq ∼ Genotype * Housing + (1| Cohort), followed by post hoc pairwise comparison with emmeans package (pairwise ∼ Genotype * Housing); ns: non-significant. Data are represented as box plot with whiskers and data points (+: mean, line: median; whiskers: minimum and maximum values).

### miR-708-5p overexpression in the mouse hippocampus is sufficient to induce MD-associated endophenotypes

We next investigated whether stress-induced upregulation of miR-708-5p in the rodent hippocampus is causally involved in the development of MD-associated behavioral endophenotypes. Towards this end, miR-708-5p was ectopically overexpressed in the mouse hippocampus using stereotactic injection of a recombinant rAAV expressing a miR-708-5p hairpin (hp) under the control of the human synapsin (hSYN) promoter (Fig. S1A). We chose mice four our studies since we our animal facilities are not set up for experiments with rats. The hippocampus was chosen for functional manipulation since it represents a key brain structure for cognitive and emotional processing, and our previous data indicates strong effects of juvenile social isolation on hippocampal miRNA expression (Martins *et al*., 2022; Valluy *et al*., 2015) (Fig. 2F). To ensure the functionality of the overexpressing construct, we performed luciferase assay upon transfection of miR-708-5p overexpressing plasmid (hp708-p) together with a luciferase sensor harboring two perfect binding sites for miR-708-5p. Thereby, we detected a significant decrease in luciferase activity upon miR-708-5p overexpression compared to the control (hpCTL-p) (Fig. S1B), demonstrating efficient repression of the reporter gene by the overexpressed miR-708-5p. We went on to perform stereotactic surgeries on seven to eight-week-old mice, which then underwent a four-week recovery period before behavioral assessments were conducted (Fig. 3A). Our approach led to a widespread infection of both the dorsal and ventral hippocampus (Fig. 3B), as well as to a robust miR-708-5p overexpression compared to control infected mice (Fig. 3C; Fig. S1C-E). Male mice overexpressing miR-708-5p in the hippocampus exhibited a significant decrease in immobility during the tail suspension test (TST), indicative of reduced behavioral despair, while this effect was not observed in female mice (Fig. 3D-E). Interestingly, neither male nor female mice with miR-708-5p overexpression showed significant differences in saccharin preference compared to control groups, suggesting no alteration in anhedonia-like behavior (Fig. S1F-G). Furthermore, we observed a strong, although non-significant trend, for elevated anxiety-related compulsive behavior, as assessed by the marble burying test (MBT), in miR-708-5p overexpressing male mice (Fig. S1H). However, results from the elevated plus maze (EPM) did not reveal significant changes in general anxiety levels between the experimental groups (Fig S1I-L). We further evaluated cognitive functions since cognitive impairments are frequently observed in MD patients. Our results from the novel object recognition (NOR) (Fig. 3G) test revealed that miR-708-5p overexpression led to a significant impairment in object recognition memory after a 24-hour delay specifically in male mice (Fig. 3H; Fig. S2A, B). This impairment was also evident in short-term memory (5 min delay), as miR-708-5p overexpressing mice failed to discriminate between familiar and novel objects after a brief five-minute interval, this time in a sex-independent manner (Fig. 3 I, J; Fig. S2C-F). Passive avoidance, however, was intact in miR-708-5p expressing male mice (Fig. S2G). Locomotor activity, assessed through the OFT remained unchanged, ruling out locomotion as a confounding factor in our behavioral assessments (Fig. S2H-J). Together, our results provide strong evidence that hippocampal overexpression of miR-708-5p leads to distinct MD-associated behaviors, including reduced behavioral despair, elevated compulsive behavior and impaired recognition memory. This behavioural “signature” is consistent with an “anti-depressant” function of miR-708-5p, for instances during the manic-like state of BD.

**Figure 3.**
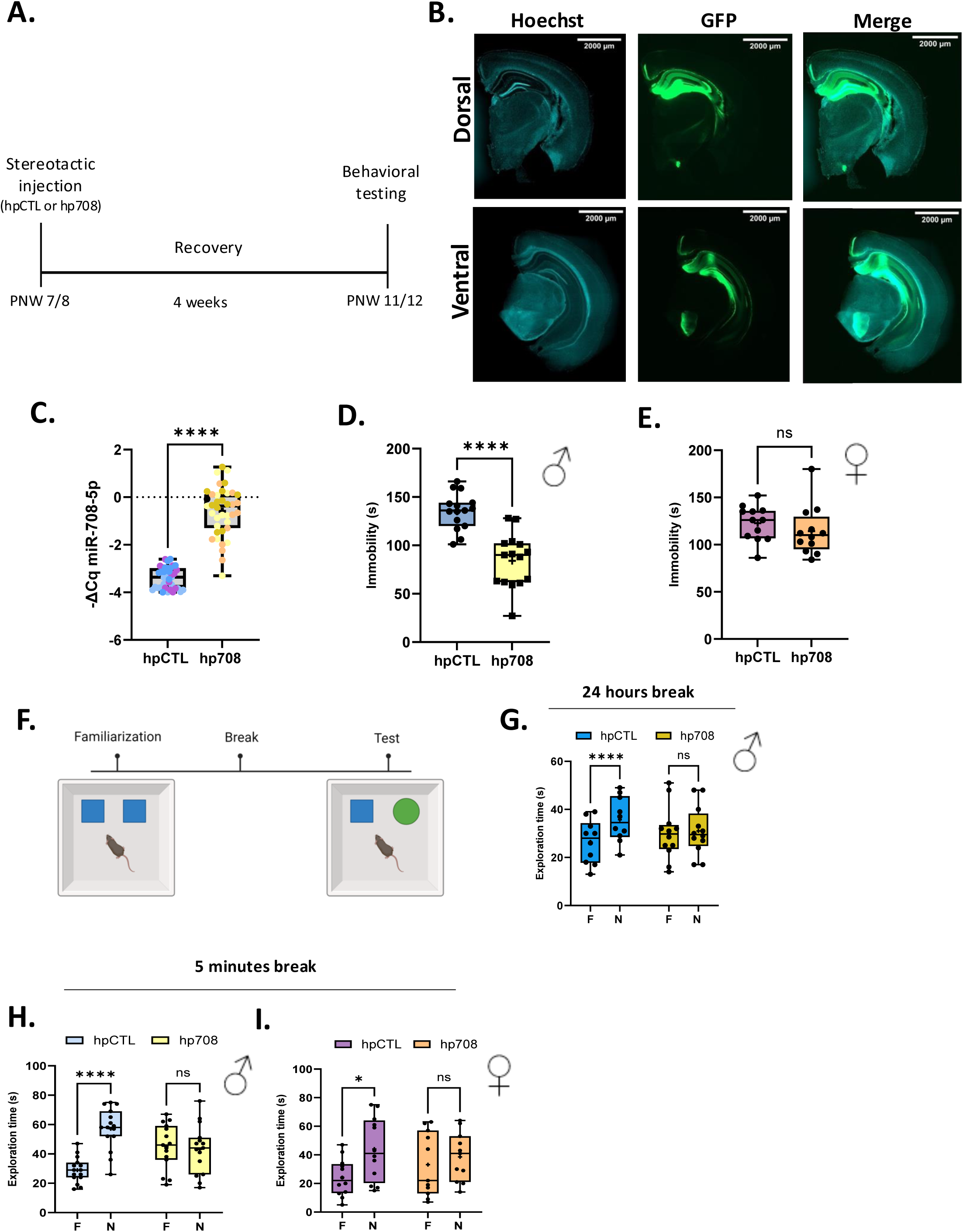
miR-708-5p overexpression in the mouse hippocampus elicits MD-associated behavioural endophenotypes. **A.** Schematic timeline of the acute rAAV stereotactic injection of control (hpCTL) or miR-708-OE (hp708) virus into the dorsal and ventral hippocampus of mice at PNW 7/8, followed by behavioral testing at PNW 11/12. **B.** Coronal brain section of mouse brains displaying intense GFP expression upon infection with hp708 virus in the dorsal (up) and ventral (down) hippocampus. Scale bar: 2000 μm. **C.** miR-708-5p qPCR analysis of total RNA isolated from hippocampus of mice that underwent behavioral testing. upon miR-708-5p overexpression. hpCTL n=39, hp708 n=39. Unpaired t-Test, ****, p<0.00001. Data are represented as box plot with whiskers and data points (+: mean, line: median; whiskers: minimum and maximum values). **D.** Tail Suspension Test. Time (s) male mice injected with the indicated rAAV (hpCTL or hp708, n=15 each) spent immobile. Data are represented as box plot with whiskers and data points (+: mean, line: median; whiskers: minimum and maximum values). Unpaired t-test, ****, p<0.0001. **E.** Tail Suspension Test. Time (s) female mice injected with the indicated rAAV (hpCTL n= 12, or hp708, n=11) spent immobile. Data are represented as box plot with whiskers and data points (+: mean, line: median; whiskers: minimum and maximum values). Unpaired t-test, ns. **F.** Schematic representation of the Novel Object Recognition (NOR) Test.mice are exposed to two familiar objects during the familiarization phase. After five minutes, they are returned to the home cage for either 24 hours or 5 minutes. They are then exposed to a novel and a familiar object during the test phase. **G.** Novel object recognition Test with 24 hours break in between familiarization and novelty testing. Time (s) male mice injected with the indicated rAAV (hpCTL n= 10, or hp708, n=12) explored either the familiar (F) or novel (N) object. Data are represented as box plot with whiskers and data points (+: mean, line: median; whiskers: minimum and maximum values). Two-way RM ANOVA: Novelty x Group, ***, p=0.0004; Novelty, ****, p<0.0001; Group, ns, p=0.8830. Šídák’s post hoc test, F vs N: hpCTL, ****, p<0.0001; hp708, ns, p=0.6735. **H.** Novel object recognition Test with 5 minutes break in between familiarization and novelty testing. Time (s) male mice injected with the indicated rAAV (hpCTL or hp708, n=15 each) explored either the familiar (F) or novel (N) object. Data are represented as box plot with whiskers and data points (+: mean, line: median; whiskers: minimum and maximum values). Two-way RM ANOVA: Novelty x Group, ****, p<0.0001; Novelty, ****, p<0.0001; Group, ns, p=0.8207. Šídák’s post hoc test, F vs N: hpCTL, ****, p<0.0001; hp708, ns, p=0.6975. **I.** Novel object recognition Test with 5 minutes break in between familiarization and novelty testing. Time (s) female mice injected with the indicated rAAV (hpCTL n= 12, or hp708, n=11) explored either the familiar (F) or novel (N) object. Data are represented as box plot with whiskers and data points (+: mean, line: median; whiskers: minimum and maximum values). Two-way RM ANOVA: Novelty x Group, *, p<0.0130; Novelty, ***, p=0.0001; Group, ns, p=0.8143. Šídák’s post hoc test, F vs N: hpCTL, ****, p<0.0001; hp708, ns, p=0.3127.

### miR-708-5p directly targets Neuronatin (Nnat), an ER-resident protein involved in calcium homeostasis

As a next step, we decided to explore the mechanisms underlying miR-708-5p dependent regulation of MD-associated behavior. miRNAs regulate gene expression through binding to mRNAs, predominantly leading to their degradation or inhibition of translation (Bartel, 2018). Therefore, we reasoned that the analysis of the hippocampal transcriptome of miR-708-5p overexpressing mice could inform us about potential miR-708-5p downstream targets mediating behavioral effects. polyA-RNA-sequencing (seq) revealed 118 differentially expressed genes (DEG) (p-value<0.01; FDR<0.001) between the hippocampus of hp-708 and hp-control infected mice (Fig. 4A). Sixteen DEGs were significantly downregulated and contained predicted miR-708-5p binding sites (Fig. 4B), making them strong candidates for direct targets of miR-708-5p. Nnat was of particular interest, since it has been previously identified as a direct target of miR-708-5p (Vatsa *et al*, 2019; Yang *et al*., 2015) and its 3’UTR contains a miR-708-5p perfect binding site (Fig. 4C). The Nnat gene encodes for a small ER-resident protein which acts as an endogenous inhibitor of the SERCA calcium pump, thereby controlling intracellular calcium homeostasis which is disturbed especially in BD (Harrison *et al*, 2019). Moreover, within the mouse brain, Nnat is primarily expressed in glutamatergic CA1 and CA3 neurons based on published single cell RNA-seq data (Fig. 4B), consistent with a functional interaction of miR-708-5p and Nnat in neurons. To further validate our RNA-seq data, we measured the levels of Nnat mRNA in the hippocampus of mice upon miR-708-5p overexpression (Fig. 4D) and detected a significant decrease compared to control infected mice (Fig. 4E). So far, our expression analysis on hippocampal tissue did not allow to distinguish between effects occurring in neuronal and non-neuronal cells. Therefore, we further extended our analysis to rat primary hippocampal neuron cultures. Like in the *in vivo* experiments, infection of rat neurons with hp708 resulted in a robust upregulation of miR-708-5p levels (Fig. 4F) and led to significant reductions in Nnat mRNA (Fig. 4G), and protein levels (Fig.4H and I), demonstrating that miR-708-5p represses endogenous Nnat expression in neurons. To prove that the downregulation of Nnat is due to a direct interaction of the miR-708-5p and Nnat mRNA via the predicted miR-708-5p binding site, we performed luciferase reporter gene assay in primary rat hippocampal neurons. Luciferase reporter genes were either fused to the wild type 3’UTR of Nnat (Nnat wt) or to a 3’UTR of Nnat containing several point mutations expected to prevent miR-708-5p binding (Nnat mt). Whereas Nnat wt was efficiently downregulated by co-transfection of hp708-p, Nnat mt was insensitive to miR-708-5p overexpression (Fig. 4J). The results obtained from these assays therefore confirm that miR-708-5p directly suppresses Nnat expression through a 3’UTR interaction. In a complementary approach, we employed a target-directed miRNA degradation (TDMD) construct (sp708-p) to reduce endogenous miR-708-5p levels (Fig. S3A-C). Here, we detected a significant increase in luciferase activity upon transfection with sp708-p compared to control for Nnat wt, but not Nnat mt expressing neurons (Fig. 4K). In summary, our results from RNA-seq, qPCR and luciferase assays establish Nnat as a direct target of miR-708-5p in rodent hippocampal neurons.

**Figure 4.**
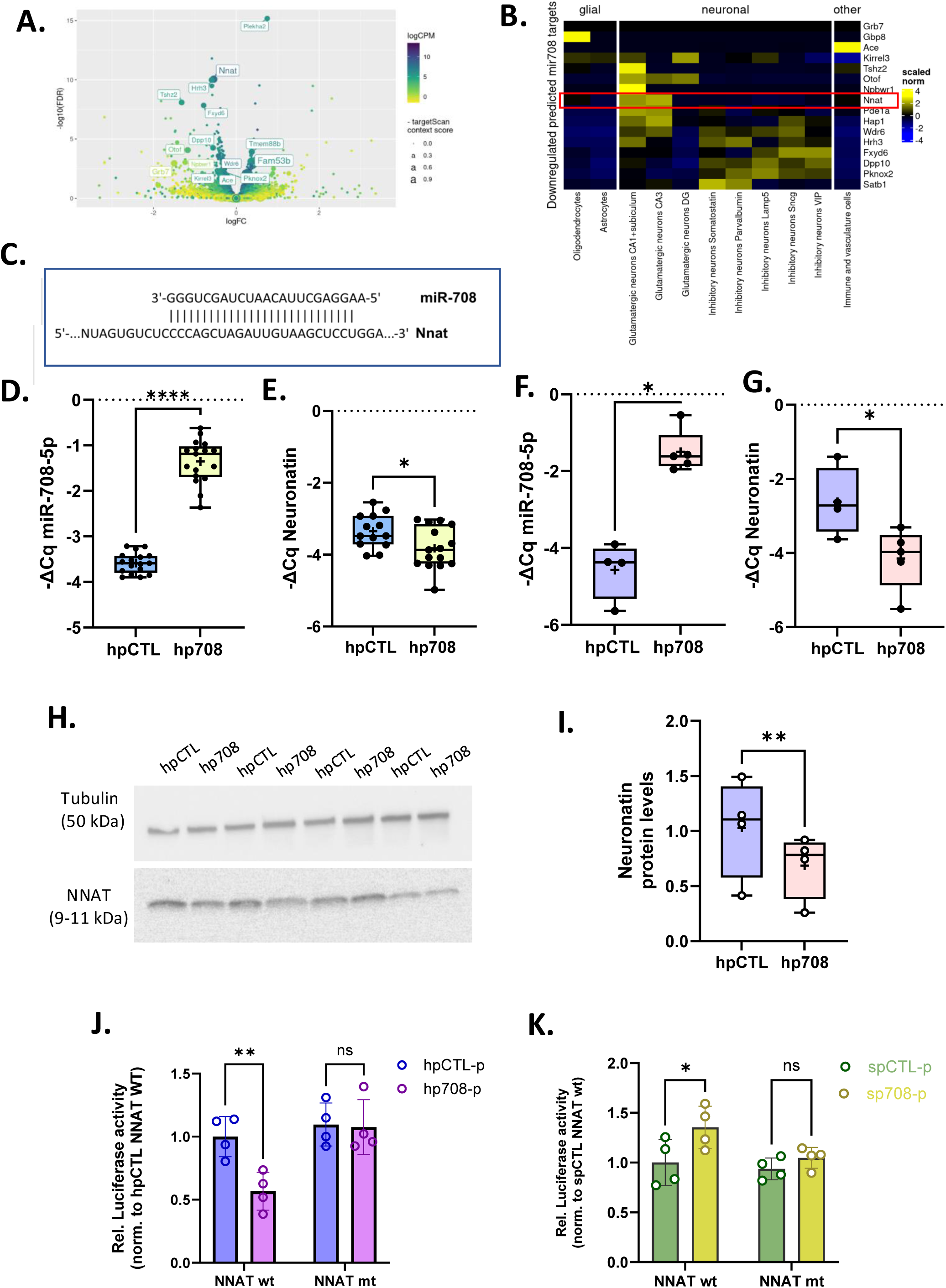
miR-708-5p directly targets Nnat, an ER-resident protein involved in calcium homeostasis. **A.** Volcano plot depicting differential gene expression in the hippocampus between hp708 and hpCTL injected mice based on polyA-RNA sequencing. **B.** Heatmap displaying miR-708-5p predicted targets which are significantly downregulated from (A), as well as their expression in different cell types based on single-cell RNA-seq data (Allen 10X+smartSeq taxonomy)(59). The red rectangle highlights Nnat. **C.** Nucleotide base pairing depicting the perfect binding of miR-708-5p with the mouse Nnat 3’-UTR. **D.** miR-708-5p qPCR analysis of total RNA isolated from mouse hippocampi upon miR-708-5p overexpression. hpCTL n=17, hp708 n=17. Unpaired t-Test, ****, p<0.00001. Data are represented as box plot with whiskers and data points (+: mean, line: median; whiskers: minimum and maximum values). **E.** Neuronatin qPCR analysis of total RNA isolated from mouse hippocampi upon miR-708-5p overexpression. hpCTL n=13, hp708 n=15. Unpaired t-Test, ****, p=0.0269. Data are represented as box plot with whiskers and data points (+: mean, line: median; whiskers: minimum and maximum values). **F.** miR-708-5p qPCR analysis of total RNA isolated from rat primary hippocampal neurons at DIV20 infected with hpCTL or hp708 for miR-708-5p overexpression. hpCTL n=4, hp708 n=5. Unpaired t-Test, *, p=0.0317. Data are represented as box plot with whiskers and data points (+: mean, line: median; whiskers: minimum and maximum values). **G.** Neuronatin qPCR analysis of total RNA isolated from rat primary hippocampal neurons at DIV20 infected with hpCTL or hp708 for miR-708-5p overexpression. hpCTL n=4, hp708 n=5. Unpaired t-Test, *, p=0.0159. Data are represented as box plot with whiskers and data points (+: mean, line: median; whiskers: minimum and maximum values). **H.** Representative Western blot image of Nnat (lower panel) and Tubulin (upper panel) protein expression levels in hippocampal neurons (DIV20) that were infected with hpCTL or hp708 for miR-708-5p overexpression on DIV2. Tubulin was used as a loading control. **I.** Quantification of the relative intensity of Nnat protein levels in hippocampal neurons (DIV20) that were infected with hpCTL or hp708 for miR-708-5p overexpression on DIV2 based on (G). Tubulin was used as normalizer. Ratio-paired t-test, **, p=0.0042. n=4 per experimental group. Data are represented as box plot with whiskers and data points (+: mean, line: median; whiskers: minimum and maximum values). **J.** Relative luciferase activity of rat hippocampal neurons transfected with the indicated plasmid (Control: hpCTL-p; overexpressing miR-708-5p: hp708-p) and expressing either Nnat wild-type (wt) or miR-708-5p binding site mutant (mt) reporter genes. Data are represented as scattered dot plots with bar, mean±SD (n=4 independent experiments; Two-way ANOVA: main effect of the hairpin *p=0.0245, of the Nnat luciferase reporter **p=0.0050, and of the hairpin by Nnat luciferase reporter interaction p<0.0374. Sidak’s post hoc test: **p=0.0092). **K.** Relative luciferase activity of rat hippocampal neurons transfected with the indicated plasmid (Control: spCTL-p; knocking-down miR-708-5p: sp708-p) and expressing either Nnat wt or mt reporter genes. Data are represented as scattered dot plots with bar, mean±SD (n=4 independent experiments; Two-way ANOVA: main effect of the hairpin *p=0.0211, no main effect of the Nnat luciferase reporter, or of the hairpin by Nnat luciferase reporter interaction. Sidak’s post hoc test: *p=0.0287).

### Restoring Nnat expression in miR-708-5p overexpressing hippocampal neurons rescues BD-associated behavioral endophenotypes in mice

Building upon our identification of Nnat as a potential regulatory target of miR-708-5p in the hippocampus, we hypothesized that Nnat downregulation might underlie the observed MD-associated behavioral alterations elicited by miR-708-5p overexpression. To test this hypothesis, we engineered a construct which allows to simultaneously overexpress Nnat and miR-708-5p upon viral infection. This was achieved by inserting the coding sequence of Nnat downstream of the hSYN promoter, coupled with a P2A self-cleaving peptide; this fragment was followed by the EGFP coding sequence and the overexpressing miR-708-5p hairpin. As a control, we utilized a similar construct where the mCherry coding sequence replaced Nnat followed by EGFP and hp708 or EGFP and hpCTL (Fig 5A). rAAV obtained from these constructs was injected into the hippocampus of seven/eight weeks old male mice by stereotaxis, followed by a four-week recovery period before behavioral assessments as described in Fig. 3A. Expression analyses confirmed the upregulation of miR-708-5p and the intended modulation of Nnat levels (Fig. 5B, 5C; Fig. S4A, and Fig. 5D, Fig. S4B). Notably, the mCherry-P2A-hp708 construct led to a pronounced upregulation of miR-708-5p (Fig. 5B) and a corresponding decrease in Nnat levels, as opposed to the control and the Nnat-P2A-hp708 conditions, the latter of which restored Nnat expression to wild-type levels (Fig. 5C and Fig. S4A, B). Behavioral analyses revealed that miR-708-5p overexpression alone impaired object discrimination ability in the NOR test as expected (Fig. 5D, Fig. S4C). This was not observed in mice injected with the Nnat-P2A-hp708 construct, which displayed a restoration of exploratory behavior towards the novel object to a similar degree than control injected mice (Fig. 5D, Fig. S4C). Furthermore, as expected from our previous results, the total exploration time of the animals was not affected in any of the conditions (Fig. S4D). Similarly, the TST results indicated reduced behavioral despair in miR-708-5p overexpressing mice, an effect that was normalized upon co-expression of Nnat, again mirroring control group behaviors (Fig. 5E). Furthermore, no significant differences between experimental groups were observed in locomotor activity in the OFT (Fig. S4E-G). Taken together, our results strongly suggest that Nnat is an important downstream target in mediating the effects on MD-associated behavioral endophenotypes caused by miR-708-5p overexpression in the mouse hippocampus.

**Figure 5.**
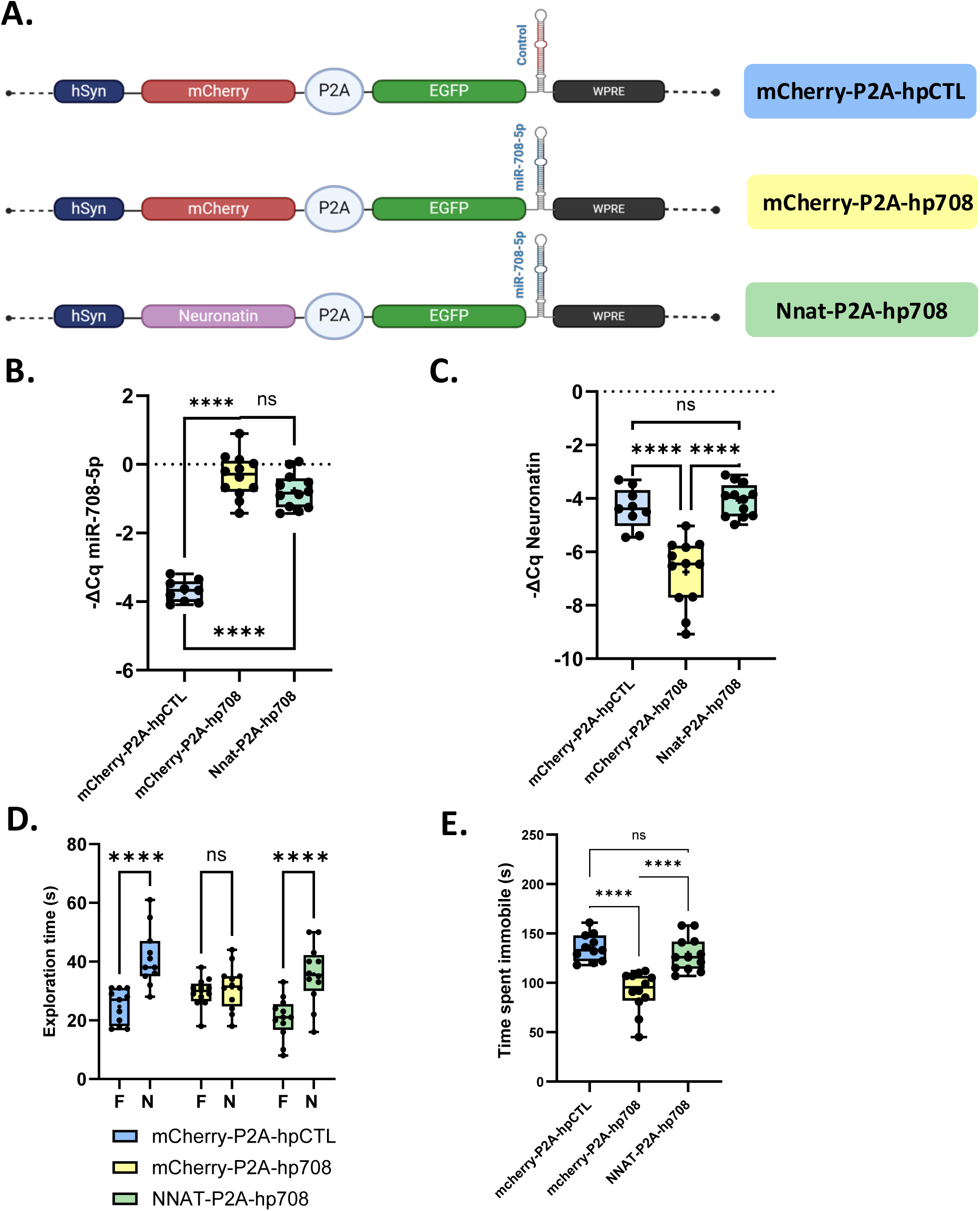
Restoring Nnat expression in miR-708-5p overexpressing hippocampal neurons rescues MD-associated behaviors in mice. **A.** Schematic representation of the three rAAV constructs used in the rescue experiment. **B.** miR-708-5p qPCR analysis of total RNA isolated from the hippocampus of male mice injected with mCherry-P2A-hpCTL (n=9), mCherry-P2A-hp708 (n12), or Nnat-P2A-hp708 (n=12) viruses. One-way ANOVA, Post hoc Tukey’s Test: mCherry-P2A-hpCTL vs mCherry-P2A-hp708: ****, p<0.0001, mCherry-P2A-hpCTL vs Nnat-P2A-hp708: ***, p<0.0001, mCherry-P2A-hp708 vs Nnat-P2A-hp708: ns, p=0.1225. Data are represented as box plot with whiskers and data points (+: mean, line: median; whiskers: minimum and maximum values). **C.** Neuronatin qPCR analysis of total RNA isolated from the hippocampus of male mice injected with mCherry-P2A-hpCTL (n=9), mCherry-P2A-hp708 (n12), or Nnat-P2A-hp708 (n=12) viruses. One-way ANOVA, Post hoc Tukey’s Test: mCherry-P2A-hpCTL vs mCherry-P2A-hp708: ***, p=0.0003, mCherry-P2A-hpCTL vs Nnat-P2A-hp708: ns, p=0.7379, mCherry-P2A-hp708 vs Nnat-P2A-hp708: ****, p<0.0001. Data are represented as box plot with whiskers and data points (+: mean, line: median; whiskers: minimum and maximum values). **D.** Novel object recognition Test with 5 minutes break in between familiarization and novelty testing. Time (s) male mice injected with the indicated rAAV (mCherry-P2A-hpCTL (n=9), mCherry-P2A-hp708 (n12), or Nnat-P2A-hp708 (n=12)) explored either the familiar (F) or novel (N) object. Data are represented as box plot with whiskers and data points (+: mean, line: median; whiskers: minimum and maximum values). Two-way RM ANOVA: Novelty x Group, ****, p<0.0001; Novelty, ****, p<0.0001; Group, ns, p=0.5765. Šídák’s post hoc test, F vs N: mCherry-P2A-hpCTL, ****, p<0.0001; mCherry-P2A-hp708, ns, p=0.6534; Nnat-P2A-hp708, ****, p<0.0001. **E.** Tail Suspension Test. Time (s) male mice injected with the indicated rAAV (mCherry-P2A-hpCTL (n=9), mCherry-P2A-hp708 (n12), or Nnat-P2A-hp708 (n=12)) spent immobile. One-way ANOVA, Post hoc Tukey’s Test: mCherry-P2A-hpCTL vs mCherry-P2A-hp708: ****, p<0.0001, mCherry-P2A-hpCTL vs Nnat-P2A-hp708: ns, p=0.9617, mCherry-P2A-hp708 vs Nnat-P2A-hp708: ****, p<0.0001. Data are represented as box plot with whiskers and data points (+: mean, line: median; whiskers: minimum and maximum values).

### miR-708-5p levels negatively correlate with human cognitive processing and represent a potential biomarker for differential diagnosis in MDs

Our functional analysis in mice implies an important role of miR-708-5p in the regulation of MD-associated behaviours. To explore a potential link between miR-708-5p and human behaviour, we harnessed our rich dataset of neuropsychological test results present within the FOR2107 cohort (Kircher *et al*., 2019). Given our previous results from mice (Fig. 3), we were specifically interested in correlations between miR-708-5p expression levels (measured in PBMCs) and performances in assessments covering the neurocognitive domains (attention/concentration; executive function, verbal and visuo-spatial memory (see materials and methods). Interestingly, miR-708-5p expression in a combined human sample (n=162; both healthy and MD subjects of both sexes; Table 1) showed a significant negative correlation with the score from the D2 attention test (Fig. 6A), which suggests that high levels of miR-708-5p in human impair selective attention and cognitive processing. Significant negative correlations were also observed when considering males (n=90) and females (n=72) separately and were mostly driven by BD and to a lesser extent MDD patients (Fig. S5). These finding aligns well with our results from behavioural testing in mice, which revealed a negative role for miR-708-5p in novel object recognition memory.

**Figure 6:**
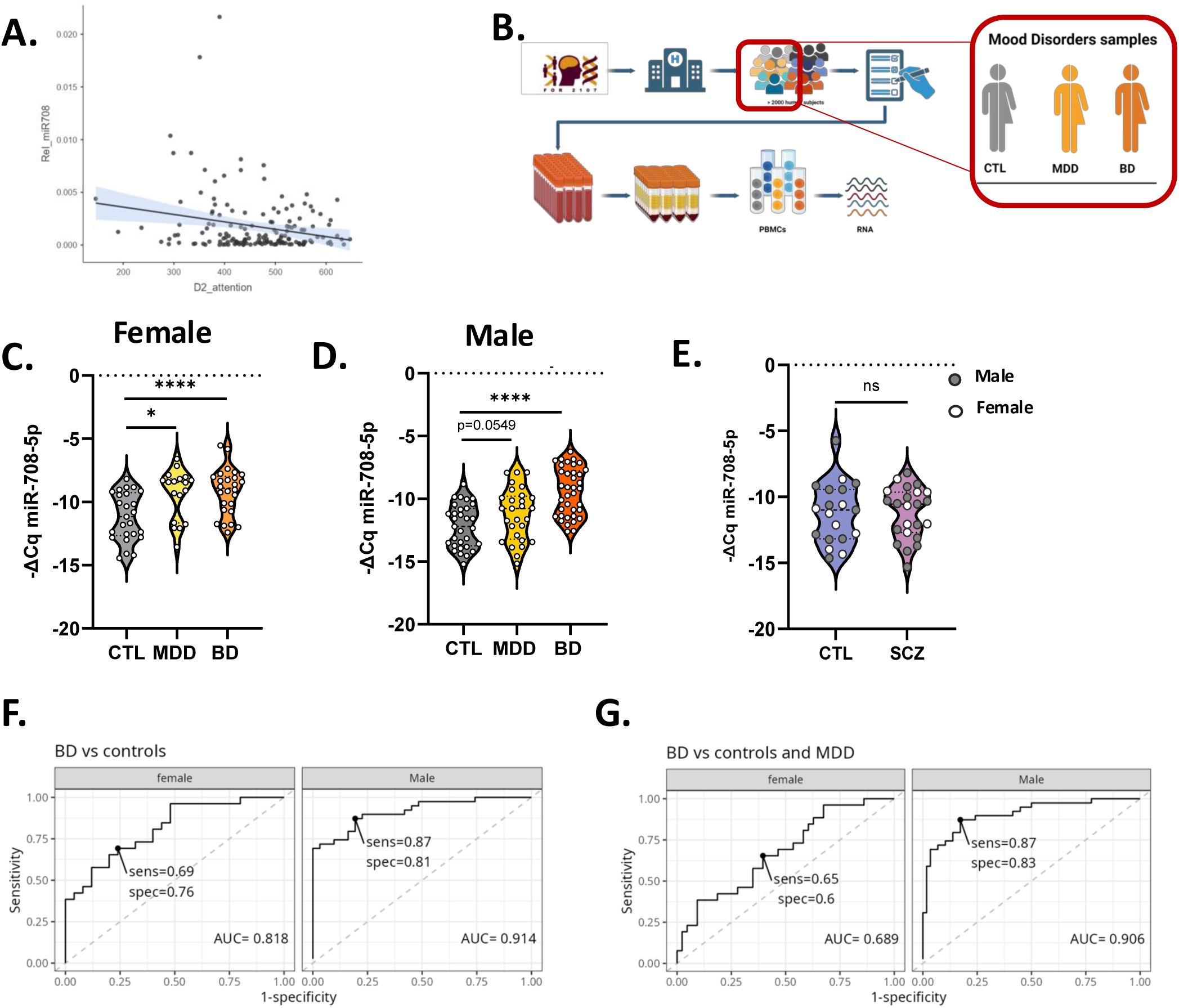
miR-708-5p levels negatively correlate with human cognitive processing and represent a potential biomarker for differential diagnosis in MDs. **A.** Pearson correlation plot of miR708 peripheral levels and attention (d2 test) in all human participants (female + male, CTL + MDD + BD, n=162), **, p=.005, r=-.223. **B.** Same as Fig. 1A, except that samples from the FOR2107 cohort were in addition collected from male and female patients that were diagnosed with Major Depressive Disorder (MDD) or Bipolar Disorder (BD) (mood disorder samples). **C.** miR-708-5p qPCR analysis of total RNA isolated from PBMCs of female patients diagnosed with BD or MDD (control: CTL= 26; MDD= 18; BD= 26). Wilcoxon rank-sum test after correction for age and antidepressant treatment; linear model of the form -DeltaCq ∼ Group + Age + Antidepressant treatment. Females: CTL vs MDD, *, p<0.05; CTL vs BD, ****, p<0.0005. **D.** miR-708-5p qPCR analysis of total RNA isolated from PBMCs of male patients diagnosed with BD or MDD (control: CTL=31; MDD=24; BD= 37. Wilcoxon rank-sum test after correction for age and antidepressant treatment; linear model of the form -DeltaCq ∼ Group + Age + Antidepressant treatment. CTL vs MDD, p=0.0549; CTL vs BD, ****, p<0.0005). Data are presented as violin plots with median, quartiles and data points. **E.** miR-708-5p qPCR analysis of total RNA isolated from PBMCs of male and female patients diagnosed with Schizophrenia (male: CTL=11; SCZ=15; female: CTL=8; SCZ=8). Two-way ANOVA: Group x Sex, ns, p=0.4904; Group, ns, p=0.2383, Sex, ns, p=0.4904. Tukey’s post hoc test, ns. Data are presented as violin plots with median, quartiles and data points. **F.** ROC curve of combined miR-708-5p and miR-499-5p expression in female (left) and male (right) subjects to discriminate BD vs control samples. The indicated thresholds are the closest point to the optimal (i.e. top-left). **G.** ROC curve of combined miR-708-5p and miR-499-5p expression in female (left) and male (right) subjects to discriminate BD vs controls and MDD samples. The indicated thresholds are the closest point to the optimal (i.e. top-left).

Finally, we assessed the potential of miR-708-5p as a diagnostic biomarker in MD, Therefore, we measured the levels of miR-708-5p in PBMCs obtained from MDD (n=42) and BD (n=63) patients (Fig. 6B; Table 1) by qPCR. miR-708-5p was significantly upregulated in female BD (n=26) and MDD (n=18) patients (Fig. 6C) compared to CTL (n=26), after correcting for age and antidepressant treatment (Fig. S6A-B), whereby the effect was more pronounced in BD compared to MDD patients. A similar picture was observed in male patients. Here, the increase was highly significant for BD subjects (Fig. 6C, after age and antidepressant treatment correction (Fig. S6C), and only trending for patients affected by MDD (Fig. 6D, Fig. S6D). Furthermore, miR-708-5p was significantly upregulated in both subtypes of BD in both female (Fig. S6E) and male (Fig. S6F) subjects. In female subjects, the only significant BD state for miR-708-5p upregulation was mania, although it only included two samples (Fig. S6G). There was also a significant miR-708-5p upregulation in depressive and hypomanic states in male subjects (Fig. S5H).

We also investigated whether miR-708-5p correlated with the Beck’s depression inventory (BDI), in MDD male and female subjects, but no significant or trending correlation could be observed (Fig. S6I and J). miR-708-5p levels showed no correlation with the Young Mania Rating Scale (YMRS), which is used to assess manic-like symptomatology in subjects affected by BD, in female BD patients (Fig. S6K). In contrast, miR-708-5p showed a trending positive correlation with YMRS in male subjects affected by BD (Fig. S6L). Taken together, our results suggest that miR-708-5p is elevated in the peripheral blood of MD patients, with a particularly strong upregulation observed for BD males.

We then asked if miR-708-5p was also differentially expressed in PBMCs obtained from patients suffering from related psychiatric disorders. For that purpose, we chose schizophrenia (SCZ), given the high genetic similarities between SCZ and BD (Craddock *et al*, 2005). We performed qPCR on PBMCs from female (n=8) and male (n=15) SCZ patients but were unable to detect any significant change in miR-708-5p expression compared to healthy control subjects (CTL; n=19) (Fig. 6E).

One of the biggest challenges in mood disorder diagnostics is to distinguish BD and MDD patients, especially when BD patients are in a depressive phase (Nierenberg *et al*, 2023; Vieta *et al*, 2018). Given our results from Fig. 6C-D, we hypothesized that miR-708-5p expression in PBMCs could be used as a diagnostic tool to classify samples into BD or control group, and BD or Control and MDD group. To do this, we performed receiver operating characteristic (ROC) curve analysis (Zweig & Campbell, 1993) and used miR-708-5p expression as predictor variable. We found that miR-708-5p alone performed better in male samples than in female samples when asked to discriminate BD patients from controls and MDD patients (Fig S7A and B), consistent with our previous results (Fig. 6C-D). We then performed the same analysis using the expression values of miR-499a-5p, which we previously found to be significantly upregulated in male BD subjects compared to healthy controls (Martins *et al*., 2022). Like miR-708-5p, miR-499a-5p performed better as a classifier in male subjects compared to female subjects (Fig S7C and D). Furthermore, we performed another ROC curve analysis considering both miRNAs together and found that the combined expression of miR-708-5p and miR-499a-5p performs better (AUC=0.914) in classifying male BD vs controls (Fig. 6F) than when the miRNAs are analyzed individually. In addition, the combination performs better (AUC=0.914) also in classifying male BD vs controls and MDD patients (AUC=0.906) (Fig. 6G).

Taken together, our results show that the combined expression of miR-708-5p and miR-499a-5p in PBMCs shows a good predictive potential to distinguish BD patients from MDD patients and healthy controls, in particular regarding males.

## Discussion

In this study, we found that miR-708-5p is upregulated in healthy human subjects at high risk to develop MDs, in genetic and environmental rat MD models, as well as in MD patients. Furthermore, peripheral miR-708-5p expression was negatively correlated with cognitive processing. Mimicking miR-708-5p overexpression in the mouse hippocampus was sufficient to elicit MD-associated behavioural alterations, namely memory impairments, increased compulsive anxiety and reduced depression-like behavior. Finally, peripheral levels of miR-708-5p, together with the previously characterized BD-associated miR-499-5p, efficiently discriminated male BD from MDD patients and healthy subjects, emphasizing its utility as a diagnostic biomarker in MDs.

### miR-708-5p regulation by genetic and environmental risk factors

MDs are highly heritable psychiatric disorders. Genetic factors seem to explain about 35–45% of variance in the etiology of MDD and 65–70% of variance for BD (Coleman *et al*, 2020; Polderman *et al*, 2015). However, genetics alone cannot explain the degree at which MD is inherited in families, suggesting that environmental factors need to be considered as well. Childhood maltreatment is the most studied environmental stressor in the context of MDs (Aas *et al*., 2020; Nemeroff, 2016). Interestingly miR-708-5p was upregulated in the juvenile social isolation rat model, an animal model for childhood maltreatment (Seffer *et al*., 2015). Consistent with this result, a recent study reported miR-708-5p upregulation in the PFC upon seven days of chronic social defeat in vulnerable rats compared to resilient animals (Chen *et al*, 2015). Moreover, miR-708-5p levels are elevated in *Cacna1c^+/-^* rats, a well-established genetic model of psychiatric disorders (Bhat *et al*., 2012), even without environmental stressors. Taken together, these results suggest that genetic and environmental risk might impinge on common pathways triggering miR-708-5p expression. Furthermore, studies in the context of ovarian cancer revealed the presence of a glucocorticoid response element upstream of *ODZ4* (Lin *et al*., 2015) , suggesting that the expression of the host gene and of miR-708-5p might be responsive to glucocorticoids. Moreover, studies in the context of ER stress suggest that miR-708-5p expression is induced by CHOP, a transcription factor involved in the Unfolded Protein Response (UPR) upon ER stress (Behrman et al., 2011). Furthermore, metformin-induced upregulation of miR-708-5p elicits Nnat downregulation in prostate cancer, leading to the expression of ER-Stress-mediators. Finally, bisphenol A, an ubiquitous endocrine disruptive chemical, induces miR-708-5p in hypothalamic neurons in a CHOP-dependent manner, accompanied by Nnat downregulation (Nierenberg et al., 2023). Thus, we speculate that increased ER stress, which has been implicated in BD, is a critical mediator of miR-708-5p upregulation in response to genetic and environmental stress in neurons. In the future, miR-708-5p manipulation in the context of a GxE rodent model should help to test this hypothesis.

### miR-708-5p function in MD-associated behaviors

Modelling the complex symptomatology of MD in animal models is challenging and has so far only been partially achieved in mice in the context of miRNA manipulation. For example, the acute manipulation of miR-124 in the PFC of mice led to impaired social behavior and locomotor disturbances upon injection of psychostimulants (Namkung *et al*, 2023). The hippocampal overexpression of miR-499-5p in *Cacna1c^+/-^* rats provoked memory impairments, recapitulating the cognitive deficits encountered in MD (Martins *et al*., 2022) which include difficulties in attention, memory and executive functions present during the different mood phases and during remission (Huang *et al*, 2023). Accordingly, we found that in humans miR-708-5p peripheral expression is negatively correlated with D2 scoring in the attention test (Camelo *et al*, 2017). Moreover, we found that both female and male mice overexpressing miR-708-5p in the hippocampus showed memory impairments. Furthermore, male, but not female, mice showed significantly decreased immobility in the TST. This test has been historically used to test antidepressant effects on mice but is also employed in mouse models of manic behavior. For example, BD mouse models of mania (Ankyrin-G conditional knockout, Clock19 deficient mice) show reduced immobility during TST (Zhu *et al*, 2017) and the FST (Roybal *et al*, 2007), suggesting that the reduced behavioral despair in our model could reflect a function of miR-708-5p in manic-like behavior. This is further supported by increased (although non-significant) marble burying, indicative of a compulsive response to anxiogenic stimuli. However, in our study, the antidepressant-like behavior was neither accompanied by increased exploratory behavior in the OFT, nor increased risk-taking during EPM testing, both of which are usually observed in mouse models of mania. We consider two possible explanations for this discrepancy. First, contrary to most of the models reported in the literature, our animal model is characterized by an acute and selective overexpression of miR-708-5p in the hippocampus. Therefore, we might expect to observe endophenotypes dependent on the hippocampus (e.g., antidepressant-like behavior, cognition), but not those related to other brain regions, such as the amygdala, prefrontal cortex, or cerebellum (e.g., risk-taking behavior, hyperactivity). Alternatively, the miR-708-5p/Nnat pathway might selectively control specific aspects of manic- and depression-like behavior. To distinguish between these possibilities, it will be important to study the role of miR-708-5p overexpression in other brain areas relevant for MD-associated behaviors in the future.

### Molecular and cellular mechanisms downstream of the miR-708-5p/Nnat interaction

Dysregulated calcium homeostasis has been previously implicated in the pathophysiology of MD, with a special emphasis on BD (Harrison *et al*., 2019). Our results from this and a previous study (Martins *et al*., 2022) are consistent with disrupted calcium flux via L-type calcium channels, e.g., due to *Cacna1c* mutation, as a major underlying cause. However, ER calcium dynamics emerges as another major player. For example, store operated calcium entry (SOCE) is dysregulated in BD-induced pluripotent stem cells, leading to earlier neuronal differentiation and abnormal neurite outgrowth (Hewitt *et al*, 2023). Furthermore, altered expression of the miR-708-5p target Nnat has been associated with defective intracellular and ER calcium levels (Sharma *et al*, 2013; Vatsa *et al*., 2019; Zou *et al*, 2023). Nnat is a small ER membrane protein which acts as an antagonist of the sarco-endoplasmic reticulum calcium ATPase (SERCA) pump, thereby interfering with calcium re-uptake into the ER (Braun *et al*, 2021). Consistently, it has been reported that miR-708-5p-mediated downregulation of Nnat causes lower basal cytoplasmic calcium levels (Vatsa *et al*., 2019). On the other hand, excessive ER calcium re-uptake could result in ER calcium overload, which in turn leads to calcium leakage to the cytoplasm and mitochondria (Daverkausen-Fischer & Prols, 2022), with negative consequences for calcium signaling and cell health, as exemplified in Alzheimer’s disease (Bezprozvanny & Mattson, 2008). Clearly, further studies are warranted on the impact of aberrant miR-708-5p/Nnat signaling on intraneuronal calcium homeostasis in the context of MD.

### miR-708-5p as a potential biomarker in BD

Our findings indicate an upregulation of miR-708-5p in PBMCs of females at risk of MDs and in patients with MDD and BD. A previous study reported a downregulated expression of miR-708-5p in leukocytes of women in depressive state compared to women in remission (Banach *et al*, 2017). These results are in line with our detected expression difference between female euthymic patients and depressive patients. For male samples, on the other hand, our data suggest a role of miR-708-5p specifically in the manic phase of the disorder, as indicated from the YMRS correlation and the behavioral characterization. Although SCZ and BD were reported to have the highest genetic correlation among psychiatric disorders (Craddock *et al*., 2005), we found that miR-708-5p expression was unchanged in peripheral samples of patients diagnosed with SCZ. This suggests a rather specific expression pattern of miR-708-5p in the spectrum of MDs. Within MDs, miR-708-5p upregulation was most pronounced in BD. This is impressively illustrated by our ROC curve analysis, which shows that miR-708-5p expression, when combined with miR-499-5p, effectively distinguishes BD patients not only from healthy controls, but also from MDD patients. Although miR-708-5p is significantly upregulated in both male and female BD patient samples, the degree of increase is higher in males compared to females (8.8-fold vs. 4.23-fold change). Moreover, based on our ROC curve analysis, miR-708-5p expression discriminated male BD patients better from control and MDD compared to female patients. Together, these observations suggest a potential sex-specific role of miR-708-5p in BD, which also aligns with our results from mouse behavior. Sex differences in BD manifest across various aspects, ranging from clinical symptoms to the progression of the disorder. Males typically encounter their first manic episode at a younger age compared to females, who are more likely to experience a depressive episode at the onset of BD (Kennedy *et al*, 2005). In contrast, females experience a higher incidence of depressive episodes and hypomania, leading to a more frequent diagnosis of BD type II (Diflorio & Jones, 2010). Females are also more susceptible to mixed episodes (Arnold *et al*, 2000), rapid cycling of mood phases (Tondo & Baldessarini, 1998), and show a greater likelihood of attempting suicide (Clements *et al*, 2013). Thus, it is tempting to speculate that miR-708-5p might play an important role in male-specific aspects of BD, e.g., the development of more common and intense manic episodes. In this regard, miRNAs have been previously implicated in the sexually dimorphic control of of circadian, cholinergic, and neurokine pathways in BD (Lobentanzer *et al*, 2019).

Taken together, we propose that miR-708-5p represents a promising candidate for the development of a biomarker that helps to stratify MD patients based on disease entity, specific phases of the disease and sex, which should greatly help with diagnosis and therapy.

## Acknowledgements

We greatly acknowledge the technical support in the preparation of primary hippocampal cultures by Cristina Furler, Tatjana Wüst and Dr. Roberto Fiore. We also thank Dr. Brunno Rocha Levone for support in behavioral testing and inputs on the manuscript. We also thank Darren Kelly for support in designing the TDMD construct, and David Colameo and Emanuel Sonder for support in the statistical analysis. RNA sequencing was performed at the Functional Genomics Center Zurich (FGCZ) of the University Zurich and ETH Zurich. The work in the lab of GS was supported by grants from ETH Zurich (ETH-24 18-2 Grant (NeuroSno)) and the Swiss National Science Foundation (SNSF <u>310030E_179651</u>, 32NE30_189486, 310030_205064/1). This work was further funded by the German Research Foundation (DFG grants FOR2107 KI588/14-1, and KI588/14-2, and KI588/20-1, KI588/22-1 to TK; grant FOR2107 DA1151/5-1, DA1151/5-2, DA1151/9-1, DA1151/10-1, DA1151/11-1 to UD; grant FOR2107 SCHR 1136/3-1 to GS; grant DFG WO 1732/4-1 and DFG WO 1732/4-2 to MW; grant DFG 559/14-1 and DFG 559/14-2 to RS), as well as the Fonds Wetenschappelijk Onderzoek – Vlaanderen (FWO; Research Foundation – Flanders) through a senior project to MW (G0C0522N), and the Interdisciplinary Center for Clinical Research (IZKF) of the medical faculty of Münster (grant Dan3/022/22 to UD). Biosamples and corresponding data were sampled, processed, and stored in the Marburg Biobank CBBMR.

## Author contributions

Carlotta Gilardi: Investigation; writing. Helena Martins: Investigation. Pierre-Luc Germain: Formal analysis. Fridolin Gross: Formal analysis. Alessandra Lo Bianco: Investigation. Silvia Bicker: Investigation; methodology. Ayse Özge Sungur: Investigation. Theresa M. Kisko: Investigation. Rainer K.W. Schwarting: Supervision. Frederike Stein: Data curation; formal analysis. Susanne Meinert: Data curation. Udo Dannlowski: supervision. Tilo Kircher: supervision; project administration. Markus Wöhr: supervision; project administration. Gerhard Schratt: Conceptualization; supervision; writing.

## Data availability

RNA sequencing data has been deposited to Gene Expression Omnibus (GEO, small RNA-sequencing Fig. 1B-C, accession numbers: GSE261287; RNA-sequencing Fig. 4A, accession numbers: GSE261288).

