## Supplementary Materials (Methods, Tables, Figures) for "Early life stress-induced miR-708-5p regulates mood disorder-associated behavioural phenotypes in mice and is a potential diagnostic biomarker for bipolar disorder"

### Supplementary Materials and Methods

#### Constructs cloning

##### Overexpressing hairpins

Briefly, the mature miR-708 duplex or a control scrambled sequence with the flanking elements of the miR-30a hairpin required for Drosha processing were cloned in the 3'UTR of eGFP on rAAV-hSyn-EGFP (Plasmid 114213, Addgene). The cloned plasmids are referred to as hpCTL-p and hp708-p; the viruses derived from these plasmids are referred to as hpCTL and hp708. The oligos used for this cloning are listed in Supplementary Table 2.

##### Knock-down constructs

To knockdown miR-708-5p a sponge plasmid was constructed by inserting six TDMD sites predicted to bind miR-708-5p in the 3'UTR of eGFP on rAAV-hSyn-EGFP. The TDMD prediction was done on the webtool ScanMIR (Uhttps://ethz-ins.org/scanMiR/). The oligonucleotide molecules used for the cloning were purchased on Thermo Fisher. The cloned plasmids are referred to as sponge-control (spCTL-p) and sponge-miR-708-5p (sp708-p). The oligos used for this cloning are listed in Supplementary Table 2.

##### pmirGLO Nnat constructs

Nnat 3'UTR was amplified from mouse brain cDNA and cloned into a pBluesScript plasmid using the Phusion Hot Start Polymerase (Thermo Fisher). Mutation of the miR-708-5p binding site was achieved using Circular Amplification for point mutations with Phusion Hot Start Polymerase (Thermo Fisher). The wild-type and mutated 3'UTRs were then shuttled into a pmirGLO dual-luciferase expression vector (Promega). To validate hairpins and sponges, two perfect binding sites for miR-708-5p were inserted into the pmirGLO dual-luciferase expression vector (Promega). The oligos used for this cloning are listed in Supplementary Table 2.

##### Rescue Constructs

The rescue constructs were cloned as followed: for mCherry-P2A-hpCTL and mCherry-P2A-hp708, the original hairpin plasmids were double digested and an oligo molecule containing the coding sequence for mCherry-P2A purchased from Thermo Fisher was inserted; For Nnat-P2A-hp708, the same procedure with a Nnat-P2a sequence was followed.

#### Tissue collection

Mice were sacrificed by cervical dislocation and the tissue was collected on an ice-cold glass plate and subsequentially snap-frozen for RNA extraction (Trizol protocol) and gene expression analysis or for histological assessments.

One hemisphere was fixed in 4% paraformaldehyde (PFA) in saline at 4°C for 2 days and then immersed in 30% sucrose in PBS. Subsequently, the tissue was frozen in Tissue-Tek O.C.T Compound (Sakura Finetek Europe B.V., 4583) and coronally sectioned in a cryostat (50 µm thick sections). These sections were promptly transferred to a cryoprotectant solution for long-term storage (0.1M Phosphate Buffer, 0.3% ethylene glycol, 1M glucose in H2O). Coronal sections were stained with Hoechst 33342 Solution (1:2000; Thermo Fisher, 62249) for 5 minutes, mounted on glass slides (Menzel-Gläser SUPERFROST PLUS, Fisher Scientific, 12-550-15), air-dried, and mounted with Aqua Poly Mount medium (Chemie Brunschwig, POL18606-20). Fourteen-bit grayscale images of GFP and Hoechst were acquired using a widefield microscope (Axio ObserverZ1/7, Zeiss) with tile scan, 5x objective, a pixel size of 1.17 µm, and an image size dependent on the section's size.

#### Stereotactic surgeries and post-operative care

Stereotactic brain injections were conducted on 3-month-old C57BL/6JRj wild-type mice. The mice were anesthetized with 5% isoflurane in oxygen (1 L/min) and positioned on a stereotaxic frame. After anaesthesia induction and before the surgical incision, the mice were transferred to a heated plate and administered 1.5-

2% isoflurane in 200 mL/min oxygen through a mouth/nose mask. To ensure their well-being, the animals received a subcutaneous injection of 5 mg/kg Meloxicam for analgesia, and vitamin A was applied to prevent eye dryness. Subsequently, the heads were shaved, cleaned, and an incision was made. Bregma and lambda were identified, and bilateral injections were performed at specific coordinates from bregma. Each rAAV virus was injected in a volume of 1 µL at each injection site using a thin capillary, with an infusion time of one minute. After a two-minute period for virus diffusion, the capillary was slowly removed. Local analgesia was achieved by suturing the skin and applying Lidocain and Bupivacain drops (2 mg/kg each) at the wound site. The mice were given time to recover from the surgery before being returned to group housing. A second subcutaneous injection of Meloxicam (5 mg/kg in saline) was administered 12 hours post-surgery, and paracetamol was added to their drinking water for the subsequent 48 hours. Postoperative health checks were conducted over the three days following the surgery.

#### Behavioral testing

For behavioral assessments, mice were individually housed and given a 20-minute acclimatization period in a holding cage before the task. The mice were then returned to their original group housing with littermates. The equipment was cleaned between trials using a detergent solution (10 ml/L Dr. Schnell AG). The behavioral assays were administered in a sequential order, progressing from the least to the most stressful.

#### Western blot

Proteins from primary hippocampal neurons were isolated using ice cold RIPA lysis buffer (10 mM Tris, pH 7.4, 150 mM NaCl, 10 mM EDTA, 2.5 mM EGTA, 1% Triton X-100, 0.1% SDS, 1% sodium deoxycholate, 10 mM NaF, 5 mM $Na_4P_2O_7$, 0.1 mM $Na_3VO_4$, supplemented with Complete Protease Inhibitor Cocktail EDTA-free (Merck, P8340). The samples were centrifuged for 5 minutes at max speed, sonicated for 10 minutes in ice-cold bath, and centrifuged at max speed. 20ug of protein was mixed with 4xLaemmli Sample Buffer (Biorad) and samples were run on a 4–20% Mini-PROTEAN® TGX™ Precast Protein Gel (Biorad). After electrophoresis, the proteins were transferred on a nitrocellulose membrane using the Trans-blot Turbo System (Biorad). Membranes were blocked for two hours at room temperature in blocking solution (2% milk in Tris-buffered saline containing 0.1% Tween20 (TBS-T)) and incubated in primary antibody (rabbit anti-Nnat, 1:1000, Abcam ab27266; rabbit anti-Tubulin, 1:2000, Cell Signaling 2125s) solution for 48 hours at 4°C. Membranes were washed five times in blocking solution and incubated in HRP (horseradish peroxidase)-conjugated secondary antibody (1:10000) in blocking solution for 1 h. Membranes were washed five times in TBS-T, developed with the Clarity™ Western ECL Substrate (Bio-Rad) and visualized with the ChemiDocTM MP, Imaging System (BioRad).

#### RT-qPCR

Total RNA extraction from brain tissue, PBMC samples, and primary hippocampal cultures was performed using TRIzol™ Reagent (Thermo Fisher, 15596026), and total RNA was subsequently extracted following the manufacturer's guidelines. The RNA was subsequentially treated with TURBO DNase enzyme (Thermo Fisher). For miRNA detection, the RNA was reverse transcribed with TaqMan MicroRNA Reverse Transcription Kit (Thermo Fisher) and the TaqMan Universal PCR Master Mix (Thermo Fisher) was used to perform RT-qPCR, according to the manufacturer's instructions. For the analysis of mRNA, the RNA was treated with the iScript cDNA synthesis kit (Bio-Rad) and the iTaq SYBR Green Supermix with ROX (Bio-Rad) was used to perform RT-qPCR on the CFX384 Real-Time System (BioRad).

#### Statistical Analysis

Statistical tests were performed using GraphPad Prism version 10.0 for Windows (GraphPad Software, San Diego, CA, USA) or R. For data sets depicting human samples, violin plots were used. The number of independent experiments is indicated in the plots. Box plots represent median (box: two quantiles around the median; whiskers: minimum and maximum value; points superimposed on the graph: individual values). Normally distributed data were tested using two-sided-Student's t-test or ANOVA followed by post hoc Tukey test and otherwise for nonnormal data the nonparametric test Mann–Whitney U-test. Novel Object Recognition and Y-maze novelty exploration time were analyzed using Two-way RM ANOVA followed by Šídák's multiple comparisons test. The distribution of data was tested with the Shapiro-Wilk test. Correlations were calculated using the Spearman correlation coefficient with two-tailed analysis. Significant changes in the BD and MDD patient group were determined by pairwise comparison using a nonparametric Wilcoxon rank sum test. To control for the effect of additional factors, a linear model of the form $-\Delta Cq \sim Group + Sex + Age + Antidepressant\ treatment$ was used in R studio. $P < 0.05$ was considered statistically significant. The detailed parameters (n, p value, test) for the statistical assessment of the data are provided in the figure legends. For ROC curves generation, to test a combination of the two miRNAs in classifying bipolar disorder patients from both controls and MDD patients, we first scaled the deltaCq values of each miRNA (to bring them to a comparable scale), and simply summed them up for each patient in order to produce the Receiver Operating Characteristic (ROC) curves (Figure 1I). For single miRNAs, the deltaCq values themselves were used to segregate the groups. The indicated thresholds are the closest point to the optimal (top-left), and the reported areas under the curves were computed using the PRROC 1.3.1 package (55). Correlational analyses were conducted to examine the relationship between relative miR-708 expression and cognitive performance. Prior to analysis, normality of the data was assessed using Shapiro-Wilk tests. Given that all data were found to be normally distributed, bivariate Pearson correlation tests were employed. All statistical analyses were conducted using Jamovi (v2.3) and R (v4.1).

#### Supplementary Table 1

Subjects used for miR-708-5p expression analysis in healthy controls (Control), healthy subjects with a history of Childhood Maltreatment (ER), healthy control with genetic risk (GR). One-way ANOVA was performed to evaluate significant differences between groups. *S.D.* standard deviation, CTQ: Childhood Maltreatment Questionnaire.

| | Control | ER | GR | P value |
|---|---|---|---|---|
| *n* | 18 | 17 | 18 | N/A |
| Sex | Female | Female | Female | N/A |
| Age ± *S.D.* | 27.6 ± 7.1 | 34.9 ± 10.9 | 29.8 ± 9.5 | 0.11 |
| CTQ ± *S.D.* | 29.7 ± 5.0 | 47.0 ± 10.4 | 30.2 ± 4.2 | <0.0001 |

#### Supplementary Table 2

| **Name of the sequence** | **Sequence (5'->3')** | **Construct** |
|---|---|---|
| **Chimeric miR-708 hairpin A_Forward** | GTACAGCTGTTGACAGTGAGCGACAAGGAGCTT | hp708 |
| **Chimeric miR-708 hairpin A_Reverse** | GATTGTAAGCTCCTTGTCGCTCACTGTCAACAGCT | hp708 |
| **Chimeric miR-708 hairpin B_Forward** | TCTAGCTGGGTGTGAAGCCACAGATGGCCCAGCTA | hp708 |
| **Chimeric miR-708 hairpin B_Reverse** | AGTAAGCTCCTTGCTGCCTACTGCCTCGGAA | hp708 |
| **Chimeric miR-708 hairpin C_Forward** | TTACTCTAGCTGGGCCATCTGTGGCTTCACACCCA | hp708 |

| | | |
|---|---|---|
| **Chimeric miR-708 hairpin C_Reverse** | AGCTTTCCGAGGCAGTAGGCAGCAAGGAG | hp708 |
| **Chimeric control hairpin A_Forward** | TGTACAGCTGTTGACAGTGAGCGACAACCTTGTG | hpCTL |
| **Chimeric control hairpin A_Reverse** | AAGGACCACAAGGTTGTCGCTCACTGTCAACAGC | hpCTL |
| **Chimeric control hairpin B_Forward** | GTCCTTAGGTGCGTGTGAAGCCACAGATGGCGC | hpCTL |
| **Chimeric control hairpin B_Reverse** | GGTTTAGGTGCGCCATCTGTGGCTTCACACGCACCT | hpCTL |
| **Chimeric control hairpin C_Forward** | ACCTAAACCACAAGGTTGCTGCCTACTGCCTCGGA | hpCTL |
| **Chimeric control hairpin C_Reverse** | AAGCTTTCCGAGGCAGTAGGCAGCAACCTTG | hpCTL |
| **miR-708-5p six TDMD sequence** | TGTACATATAACTAGTTGCATTGTCGATTCATCCCAGCTAGATT GCAATAGCTCCTTGACTCCCAGCTAGATTGCAATAGCTCCTTC AGTCCCAGCTAGATTGCAATAGCTCCTTGACTCCCAGCTAGAT TGCAATAGCTCCTTACGTCCCAGCTAGATTGCAATAGCTCCTT GTCACCCAGCTAGATTGCAATAGCTCCTTGATCGTAGCTAAGC TTGTCGAC | sp708 |
| **Control six TDMD sequence** | TGTACATATAACTAGTTGCATTGTCGATTCATTGCTACATGGTC GGAACGGTCTGACTATCTTCGAGGTCGGTTTCTTACCGCTCA CTTAGTGGTCCGTCGGTGATGACTATCATGCTGGTCACTTTG GCGGACGTCTCTATGAGGTCTATGGTCCGGGTCATCTTGACA GGTCAGGCGTTCGTGATCGTAGCTAAGCTTGTCGAC | spCTL |
| **PBS miR708-fw** | ACTAGTCCCAGCTAGATTGTAAGCTCCTTctCCCAGCTAGATTG TAAGCTCCTT | pMIR-GLO-2xPBS miR-708-5p |
| **PBS miR708-rv** | AAGCTTAAGGAGCTTACAATCTAGCTGGGagAAGGAGCTTAC AATCTAGCTGGG | pMIR-GLO-2xPBS miR-708-5p |
| **Nnat 3'-UTR fw** | TAAGCAGAGCTCCCCCAGCTCCCAGCCct | pMiR-GLO-Nnat3'UTR-WT |

| | | |
|---|---|---|
| **feNnat 3′-UTR rv** | TAAGCAGTCGACTTTTGGTGCACCCCCACT | pMiR-GLO-Nnat3′UTR-WT |
| **Nnat 3′-UTR mutagenesis fw** | CGAGCTACATTgTACGCTGCTGGAGACAGGGACCACCTC | pMiR-GLO-Nnat3′UTR-mt |
| **Nnat 3′-UTR mutagenesis rv** | AGCAGCGTAcAATGTAGCTCGGGAGACACTACTAATGCACACTT | pMiR-GLO-Nnat3′UTR-mt |

### Supplementary Figure legends

#### Supplementary Figure 1

**A.** Schematic representation of the overexpressing hairpin of miR-708-5p placed in the 3'-UTR of EGFP.

**B.** Relative luciferase activity of rat hippocampal neurons transfected with the indicated plasmid (Control: hpCTL-p; overexpressing miR-708-5p: hp708-p) and a luciferase reporter expressing 2x Perfect Binding sites (PBS) for miR-708-5p. Data are represented as scattered dot plots with bar, mean±SD (n=3 independent experiments; Ratio paired t test, two-tailed, **, p=0.0057).

**C. , D. and E**. miR-708-5p qPCR analysis of total RNA isolated from mouse hippocampi upon miR-708-5p overexpression after behavioral characterization. Each bar corresponds to one animal. Data are represented as nested bar graphs.

**F.** Saccharin Preference Test. Cumulative Saccharin Preference (%) of male mice injected with the indicated rAAV (hpCTL or hp708, n=10 each). Data are represented as bar plot and mean±SD. Two-way RM ANOVA: Timepoint x Group, ns, p=0.6645; Timepoint, **, p=0.0067; Group, ns, p=0.1088. Šídák's post hoc test, hpCTL vs hp708: 12h, ns, p=0.6407; 24h, ns, p=0.2698; 36h, ns, p=0.3400; 48h, ns, p=0.5744.

**G.** Saccharin Preference Test. Cumulative Saccharin Preference (%) of female mice injected with the indicated rAAV (hpCTL n= 12, or hp708, n=11). Data are represented as bar plot and mean±SD. Two-way RM ANOVA: Timepoint x Group, ns, p=0.7385; Timepoint, *, p=0.0489; Group, ns, p=0.1098. Šídák's post hoc test, hpCTL vs hp708: 12h, ns, p=0.3540; 24h, ns, p=0.2635; 36h, ns, p=0.7392; 48h, ns, p=0.6978.

**H.** Marble Burying test. Number of marbles male mice injected with the indicated rAAV (hpCTL n= 11, or hp708, n=12) buried in 30 minutes. Data are represented as box plot with whiskers and data points (+: mean, line: median; whiskers: minimum and maximum values). Unpaired t-test, ns, p=0.0524.

**I. and J**. Elevated Plus Maze test. Time (s) male mice injected with the indicated rAAV (hpCTL or hp708, n=15 each) spent in closed or open arm. Data are represented as box plot with whiskers and data points (+: mean, line: median; whiskers:

minimum and maximum values). Unpaired t-test, open arms, ns, p=0.5091; closed arms, ns, p=0.9516.

**K and L.** Elevated Plus Maze test. Time (s) female mice injected with the indicated rAAV (hpCTL n= 12, or hp708, n=11) spent in closed or open arms. Data are represented as box plot with whiskers and data points (+: mean, line: median; whiskers: minimum and maximum values). Unpaired t-test, closed arms, ns, p=0.1412; open arms, ns, p=0.2364.

#### Supplementary Figure 2

**A.** Novel object recognition Test with 24 hours break in between familiarization and novelty testing. Time (s) male mice injected with the indicated rAAV (hpCTL n= 12, or hp708, n=11) explored the familiar (F) and novel (N) object. Data are represented as box plot with whiskers and data points (+: mean, line: median; whiskers: minimum and maximum values). Unpaired t-test, ns, p=0.8869

**B.** Discrimination index calculated as time spent exploring novel object / time spent exploring novel and familiar objects for female mice male mice injected with the indicated rAAV (hpCTL n= 10, or hp708, n=12). Data are represented as scattered dot plots with bar, mean±SD.

**C.** Novel object recognition Test with 5 minutes break in between familiarization and novelty testing. Time (s) male mice injected with the indicated rAAV (hpCTL or hp708, n=15 each) explored the familiar (F) and novel (N) object. Data are represented as box plot with whiskers and data points (+: mean, line: median; whiskers: minimum and maximum values). Unpaired t-test, ns, p=0.8207.

**D.** Discrimination index calculated as time spent exploring novel object / time spent exploring novel and familiar objects for male mice male mice injected with the indicated rAAV (hpCTL or hp708, n=15 each). Data are represented as scattered dot plots with bar, mean±SD.

**E.** Novel object recognition Test with 5 minutes break in between familiarization and novelty testing. Time (s) female mice injected with the indicated rAAV (hpCTL n= 12, or hp708, n=11) explored the familiar (F) and novel (N) object. Data are represented as box plot with whiskers and data points (+: mean, line: median; whiskers: minimum and maximum values). Unpaired t-test, ns, p=0.9257.

**F.** Discrimination index calculated as time spent exploring novel object / time spent exploring novel and familiar objects for female mice male mice injected with the indicated rAAV (hpCTL n= 12, or hp708, n=11). Data are represented as scattered dot plots with bar, mean±SD.

**G.** Passive Avoidance Test. Latency (s) male mice injected with the indicated rAAV (hpCTL n= 10, or hp708, n=12) to enter the dark box in Phase1 and Phase 2. Data are represented as box plot with whiskers and data points (+: mean, line: median; whiskers: minimum and maximum values). Two-way RM ANOVA: Phase x Group, ns, p=0.6815; Phase, ***, p=0.0003; Group, ns, p=0.9451. Šídák's post hoc test, Phase 1 vs Phase 2: hpCTL, *, p=0.0265; hp708, **, p=0.0036.

**H.** Open Field Test. Total Distance travelled (cm) by male mice injected with the indicated rAAV (hpCTL or hp708, n=15 each). Data are represented as XY axis with Mean±SD.

**I.** Open Field Test. Total Distance travelled (cm) by female mice injected with the indicated rAAV (hpCTL n= 12, or hp708, n=11). Data are represented as XY axis with Mean±SD.

**J.** Open Field Test. Total Distance travelled (cm) by male mice injected with the indicated rAAV (hpCTL or hp708, n=12each). Data are represented as XY axis with Mean±SD.

#### Supplementary Figure 3

**A.** Schematic representation of the construct to knock-down of miR-708-5p via six TDMD sites.

**B.** Prediction of miR-708-5p binding on the TDMD sites of the construct in C. using the ScanMiRApp ( https://ethz-ins.org/scanMiR/ ).

**C.** Relative luciferase activity of rat hippocampal neurons transfected with the indicated plasmid (Control: spCTL-p; knocking-down miR-708-5p: sp708-p) and a luciferase reporter expressing 2x Perfect Binding sites (PBS) for miR-708-5p. Data are represented as scattered dot plots with bar, mean±SD (n=3 independent experiments; Ratio paired t-test, two-tailed, *, p=0.0372.

#### Supplementary Figure 4

**A. and B**. miR-708-5p and Nnat qPCR analysis of total RNA isolated from mouse hippocampi upon miR-708-5p overexpression after behavioral characterization. Each bar corresponds to one animal. Data are represented as nested bar graphs.

**C**. Novel object recognition Test. Discrimination index calculated as time spent exploring novel object / time spent exploring novel and familiar objects for male mice injected with mCherry-P2A-hpCTL (n=9), mCherry-P2A-hp708 (n12), or Nnat-P2A-hp708 (n=12) viruses. Data are represented as scattered dot plots with bar, mean±SD.

**D.** Novel object recognition Test. Time (s) male mice injected with mCherry-P2A-hpCTL (n=9), mCherry-P2A-hp708 (n12), or Nnat-P2A-hp708 (n=12) viruses explored the familiar (F) and novel (N) object. Data are represented as box plot with whiskers and data points (+: mean, line: median; whiskers: minimum and maximum values). One-way ANOVA, mCherry-P2A-hpCTL vs mCherry-P2A-hp708, ns, p=0.9439; mCherry-P2A-hpCTL vs Nnat-P2A-hp708, ns, p=0.5747; mCherry-P2A-hp708 vs Nnat-P2A-hp708, p=0.7396.

**E.** Open Field Test. Time (s) male mice injected with mCherry-P2A-hpCTL (n=9), mCherry-P2A-hp708 (n12), or Nnat-P2A-hp708 (n=12) viruses spent exploring the center of the arena. Data are represented as box plot with whiskers and data points (+: mean, line: median; whiskers: minimum and maximum values). One-way ANOVA, mCherry-P2A-hpCTL vs mCherry-P2A-hp708, ns, p=0.8167; mCherry-P2A-hpCTL vs Nnat-P2A-hp708, ns, p>0.9999; mCherry-P2A-hp708 vs Nnat-P2A-hp708, p=0.7917.

**F.** Open Field Test. Time (s) male mice injected with mCherry-P2A-hpCTL (n=9), mCherry-P2A-hp708 (n12), or Nnat-P2A-hp708 (n=12) viruses spent exploring the periphery of the arena. Data are represented as box plot with whiskers and data points (+: mean, line: median; whiskers: minimum and maximum values). One-way ANOVA, mCherry-P2A-hpCTL vs mCherry-P2A-hp708, ns, p=0.8171; mCherry-P2A-hpCTL vs Nnat-P2A-hp708, ns, p>0.9999; mCherry-P2A-hp708 vs Nnat-P2A-hp708, p=0.7922.

**G.** Open Field Test. Total Distance travelled (cm) by male mice injected with mCherry P2A-hpCTL (n=9), mCherry-P2A-hp708 (n12), or Nnat-P2A-hp708 (n=12) viruses. Data are represented as XY axis with Mean$\pm SD$.

#### Supplementary Figure 5

**A.** Pearson correlation plot of miR708 peripheral levels and attention (d2 test) in all male participants (healthy control (HC) + MDD + BD, n=92), *, p=.047, r=-.221.

**B.** Pearson correlation plot of miR708 peripheral levels and attention (d2 test) in all male participants (healthy control (HC) + MDD + BD, n=92), split by disorder group.

**C.** Pearson correlation plot of miR708 peripheral levels and attention (d2 test) in all female participants (healthy control (HC) + MDD + BD, n=70), ***, p=.006, r=-.324.

**D.** Pearson correlation plot of miR708 peripheral levels and attention (d2 test) in all female participants (healthy control (HC) + MDD + BD, n=70), split by disorder group.

#### Supplementary Figure 6

**A.** miR-708-5p qPCR analysis of total RNA isolated from PBMCs of female subjects (control, n=26; BD, n= 19; BD+AD, n=7). Kruskal Wallis Test, Post hoc Dunn's Test: Control vs BD: **, p=0.0091, Control vs BD+AD: **, p=0.0089, BD vs BD+AD: ns, p>0.9999. Data are presented as violin plots with median, quartiles and data points.

**B.** miR-708-5p qPCR analysis of total RNA isolated from PBMCs of female subjects (control, n=26; MDD, n= 2; MDD+AD, n=16). Kruskal Wallis Test, Post hoc Dunn's: Control vs MDD: ns, p=0.0532, Control vs MDD+AD: ns, p=0.0295, MDD vs MDD+AD: ns, p=0.6661. Data are presented as violin plots with median, quartiles and data points.

**C.** miR-708-5p qPCR analysis of total RNA isolated from PBMCs of male subjects (control, n=31; BD, n= 20; BD+AD, n=17). Kruskal Wallis Test, Post hoc Dunn's Test: Control vs BD: ***, p=0.0004, Control vs BD+AD: ***, p=0.0002, BD vs BD+AD: ns, p>0.9999. Data are presented as violin plots with median, quartiles and data points.

**D.** miR-708-5p qPCR analysis of total RNA isolated from PBMCs of male subjects (control, n=31; MDD, n= 11; MDD+AD, n=17). One-way ANOVA, Post hoc Tukey's test: Control vs MDD: ns, p=0.0626, Control vs MDD+AD: ns, p=0.5181, MDD vs MDD+AD: ns, p=0.4368. Data are presented as violin plots with median, quartiles and data points.

**E.** miR-708-5p qPCR analysis of total RNA isolated from PBMCs of female subjects (control, n=26; BDI, n= 10; BD II, n=14). Kruskal Wallis Test, Post hoc Dunn's: Control vs BD I: *p=0.0253, Control vs BD II: *p=0.0196, BD I and BD II: ns, p>0.9999. Data are presented as violin plots with median, quartiles and data points.

**F.** miR-708-5p qPCR analysis of total RNA isolated from PBMCs of male subjects (control, n=31; BDI, n= 16; BD II, n=23). One-way ANOVA, Post hoc Tukey's test: Control vs BD I: ***p=0.0005, Control vs BD II: ****p=0.0001, BD I and BD II: ns, p=0.2690. Data are presented as violin plots with median, quartiles and data points.

**G.** miR-708-5p qPCR analysis of total RNA isolated from PBMCs of female subjects (control, n= 26) and BD subjects in different mood states (depressive, n=7; euthymic, n= 7; hypomanic, n=25; manic, n=2; mixed, n=1). Kruskal Wallis Test, Post hoc Dunn's: Control vs. Manic, *p=0.0300. Data are presented as violin plots with median, quartiles and data points.

**H.** miR-708-5p qPCR analysis of total RNA isolated from PBMCs of male subjects (control, n= 31) and BD subjects in different mood states (depressive, n=12; euthymic, n= 5; hypomanic, n=5; manic, n=2; mixed, n=1). One-way ANOVA,

Post hoc Dunnett's test: Control vs. Depressive, ***p=0.0002; Control vs. Hypomanic, *p=0.0490. Data are presented as violin plots with median, quartiles and data points.

**I.** Correlation plot between the peripheral levels of miR-708-5p in female MDD samples (n=18) and Beck's Depression Inventory. Spearman correlation coefficient with two-tailed analysis. Data are presented as XY tables.

**J.** Correlation. plot between the peripheral levels of miR-708-5p in male MDD samples (n=28) and Beck's Depression Inventory. Spearman correlation coefficient with two-tailed analysis. Data are presented as XY tables.

**K.** Correlation plot between the peripheral levels of miR-708-5p in female BD samples (n=26) and Young Mania Rating Scale. Spearman correlation coefficient with two-tailed analysis. Data are presented as XY tables.

**L.** Correlation plot between the peripheral levels of miR-708-5p in male BD samples (n=37) and Young Mania Rating Scale. Spearman correlation coefficient with two-tailed analysis. Data are presented as XY tables.

#### Supplementary Figure 7

**A. and B**. ROC curves of miR-708-5p expression in female (left) and male (right) subjects to discriminate BD vs controls and MDD samples (top) and BD vs controls (down). The indicated thresholds are the closest point to the optimal (i.e. top-left).

**C And D**. ROC curves of miR-499-5p expression in female (left) and male (right) subjects to discriminate BD vs controls and MDD samples (top) and BD vs controls (down). The indicated thresholds are the closest point to the optimal (i.e. top-left).

**Suppl. Fig. 1**

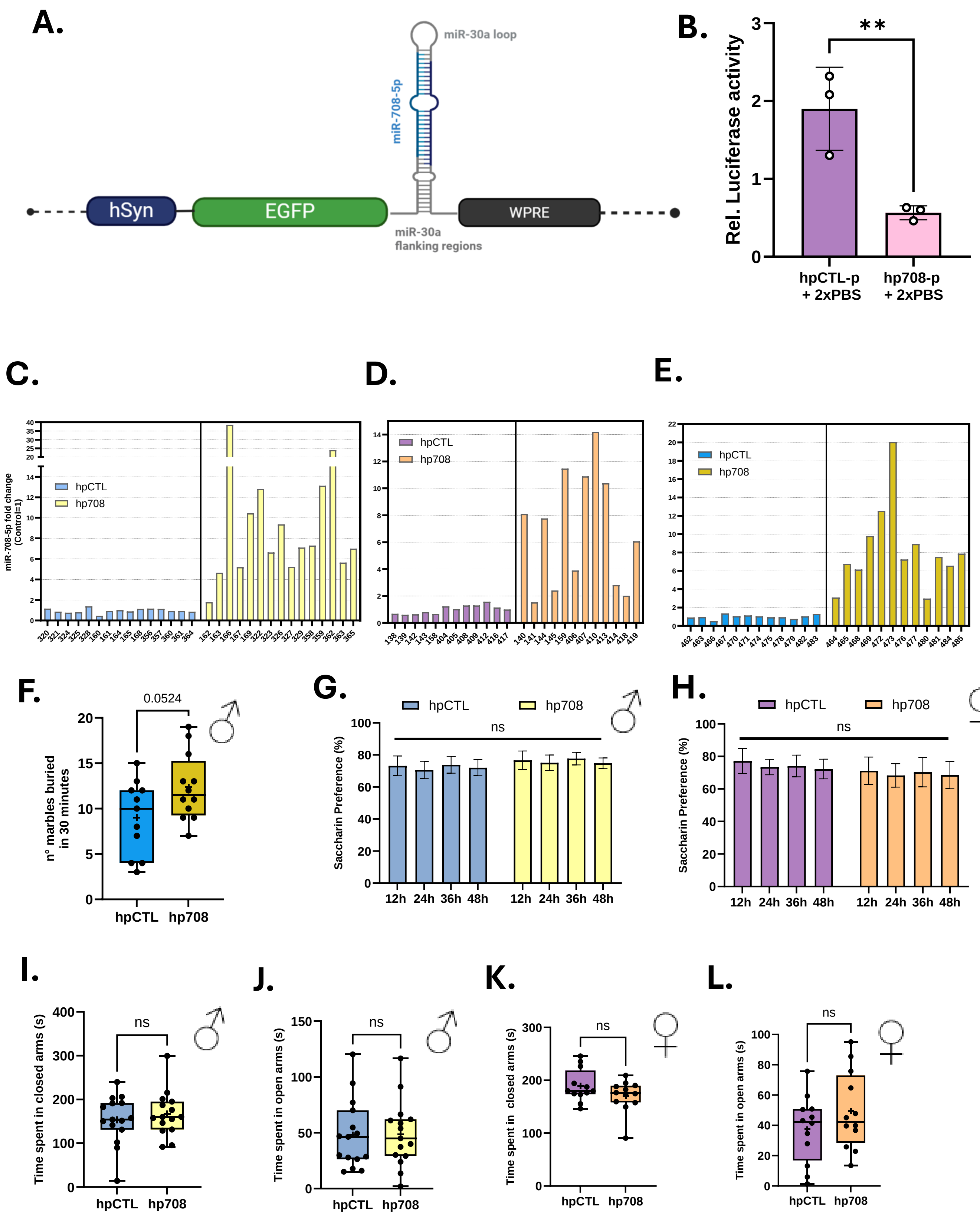
A.
miR-30a loop
miR-708-5p
hSyn
EGFP
WPRE
miR-30a
flanking regions
B.
Rel. Luciferase activity
**
hpCTL-p
+ 2xPBS
hp708-p
+ 2xPBS
C.
miR-708-5p fold change
(Control=1)
hpCTL
hp708
D.
hpCTL
hp708
E.
hpCTL
hp708
F.
0.0524
n° marbles buried
in 30 minutes
hpCTL
hp708
G.
hpCTL
hp708
ns
Saccharin Preference (%)
12h
24h
36h
48h
H.
hpCTL
hp708
ns
Saccharin Preference (%)
12h
24h
36h
48h
I.
ns
Time spent in closed arms (s)
hpCTL
hp708
J.
ns
Time spent in open arms (s)
hpCTL
hp708
K.
ns
Time spent in closed arms (s)
hpCTL
hp708
L.
ns
Time spent in open arms (s)
hpCTL
hp708

**Suppl. Fig. 2**

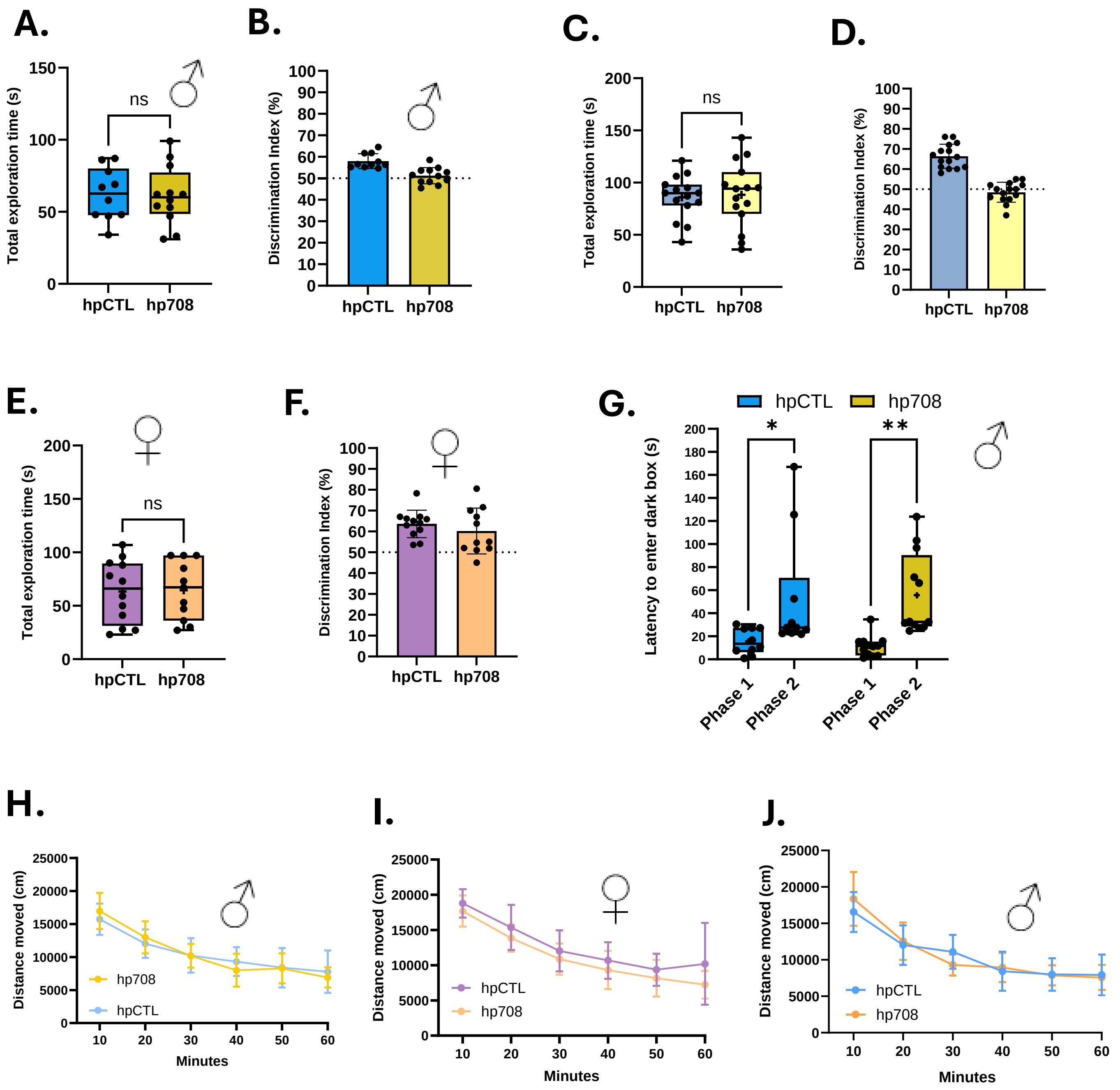
A.
Total exploration time (s)
ns
hpCTL
hp708
B.
Discrimination Index (%)
hpCTL
hp708
C.
Total exploration time (s)
ns
hpCTL
hp708
D.
Discrimination Index (%)
hpCTL
hp708
E.
Total exploration time (s)
ns
hpCTL
hp708
F.
Discrimination Index (%)
hpCTL
hp708
G.
hpCTL
hp708
Latency to enter dark box (s)
*
**
Phase 1
Phase 2
Phase 1
Phase 2
H.
Distance moved (cm)
hp708
hpCTL
Minutes
I.
Distance moved (cm)
hpCTL
hp708
Minutes
J.
Distance moved (cm)
hpCTL
hp708
Minutes

Supplementary Fig.3

A.

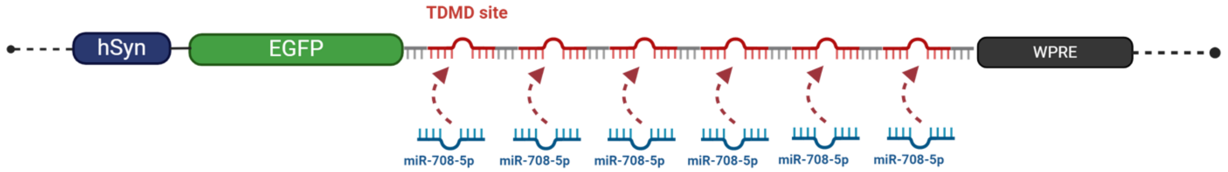


B.

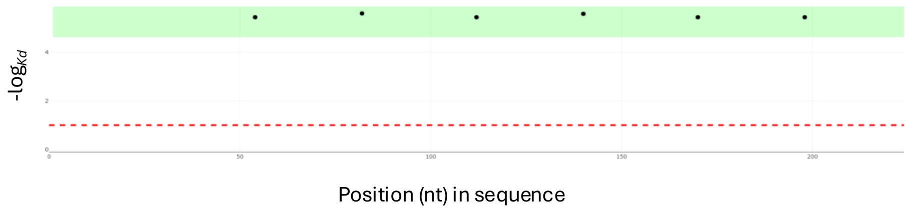


C.

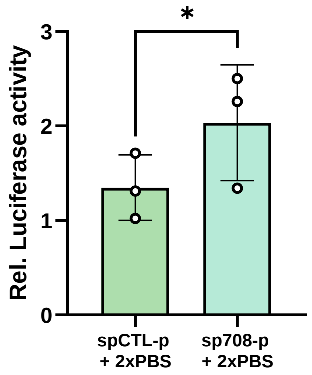

**Suppl. Fig. 4**

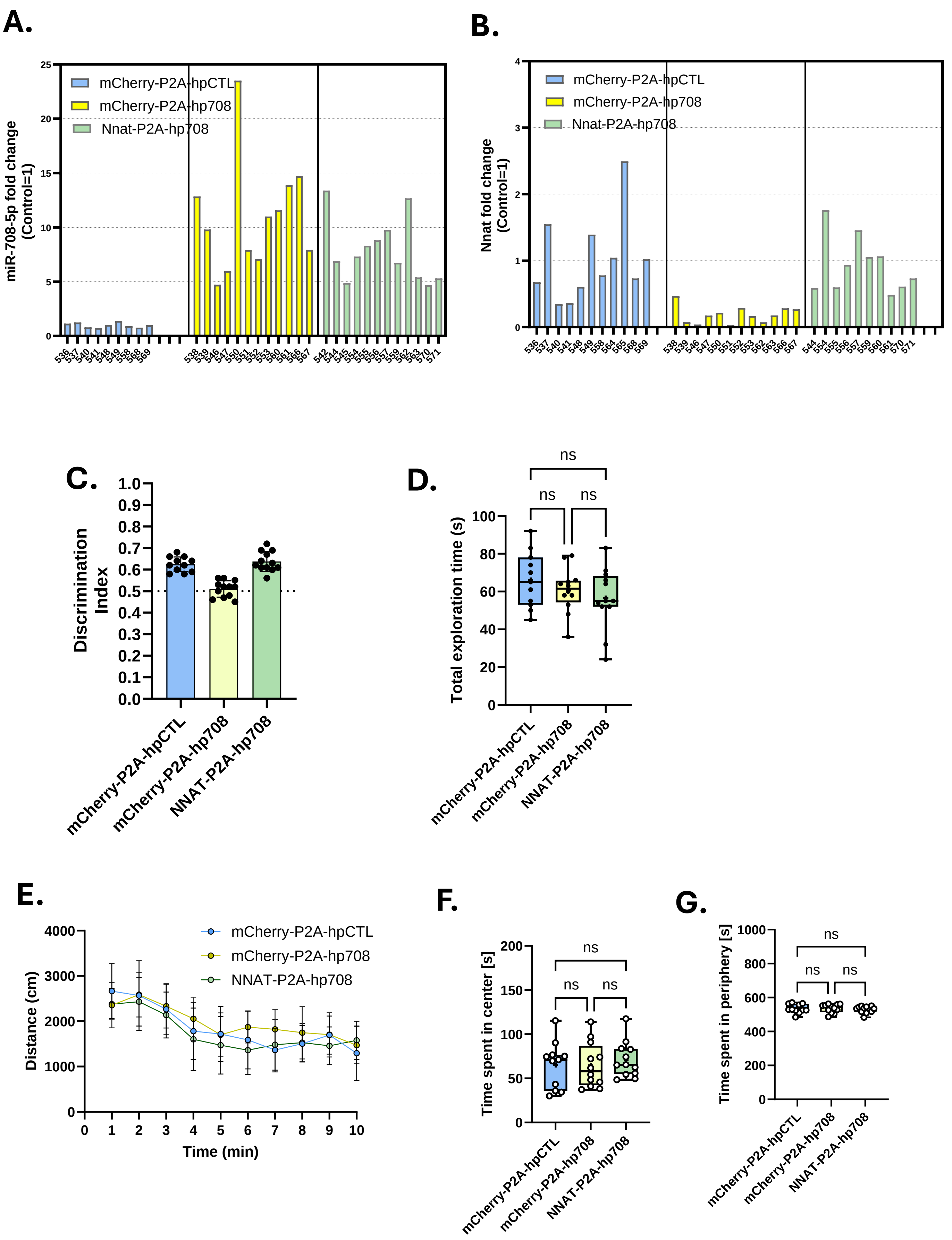
A.
miR-708-5p fold change (Control=1)
mCherry-P2A-hpCTL
mCherry-P2A-hp708
Nnat-P2A-hp708
0
5
10
15
20
25
536 537 540 541 548 549 558 568 569
538 539 546 547 550 551 552 553 560 561 566 567
542 544 545 554 555 556 557 559 562 563 570 571
B.
Nnat fold change (Control=1)
mCherry-P2A-hpCTL
mCherry-P2A-hp708
Nnat-P2A-hp708
0
1
2
3
4
536 537 540 541 548 549 558 564 565 568 569
538 539 546 547 550 551 552 553 562 563 566 567
544 554 555 556 557 559 560 561 570 571
C.
Discrimination Index
0.0 0.1 0.2 0.3 0.4 0.5 0.6 0.7 0.8 0.9 1.0
mCherry-P2A-hpCTL
mCherry-P2A-hp708
NNAT-P2A-hp708
D.
Total exploration time (s)
0 20 40 60 80 100
ns
ns
ns
mCherry-P2A-hpCTL
mCherry-P2A-hp708
NNAT-P2A-hp708
E.
Distance (cm)
0 1000 2000 3000 4000
Time (min)
0 1 2 3 4 5 6 7 8 9 10
mCherry-P2A-hpCTL
mCherry-P2A-hp708
NNAT-P2A-hp708
F.
Time spent in center [s]
0 50 100 150 200
ns
ns
ns
mCherry-P2A-hpCTL
mCherry-P2A-hp708
NNAT-P2A-hp708
G.
Time spent in periphery [s]
0 200 400 600 800 1000
ns
ns
ns
mCherry-P2A-hpCTL
mCherry-P2A-hp708
NNAT-P2A-hp708

**Suppl. Fig. 5**

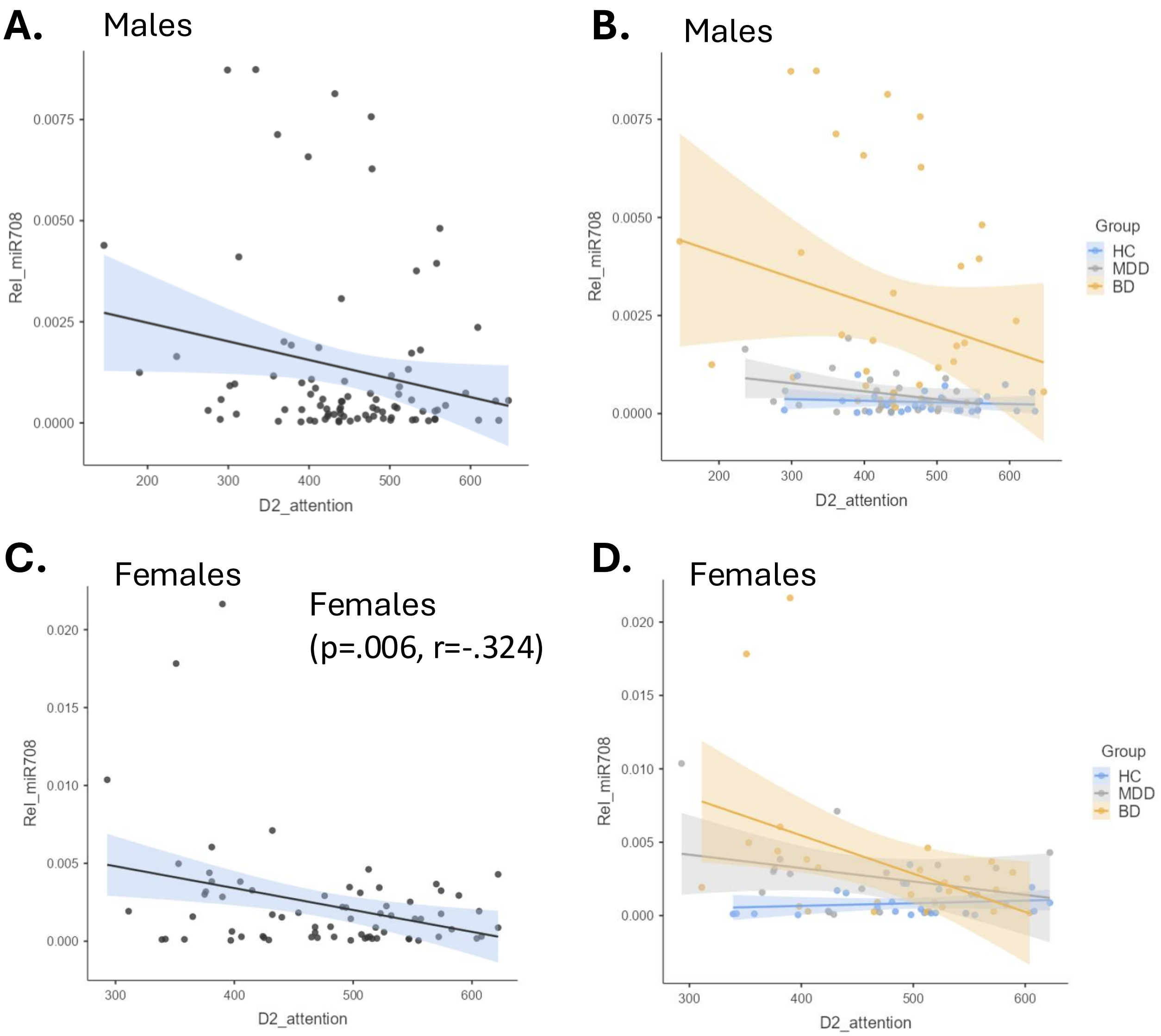
A.
Males
Rel_miR708
0.0075
0.0050
0.0025
0.0000
200
300
400
500
600
D2_attention
B.
Males
Rel_miR708
0.0075
0.0050
0.0025
0.0000
200
300
400
500
600
D2_attention
Group
HC
MDD
BD
C.
Females
Females
(p=.006, r=-.324)
Rel_miR708
0.020
0.015
0.010
0.005
0.000
300
400
500
600
D2_attention
D.
Females
Rel_miR708
0.020
0.015
0.010
0.005
0.000
300
400
500
600
D2_attention
Group
HC
MDD
BD

Suppl. Fig. 6

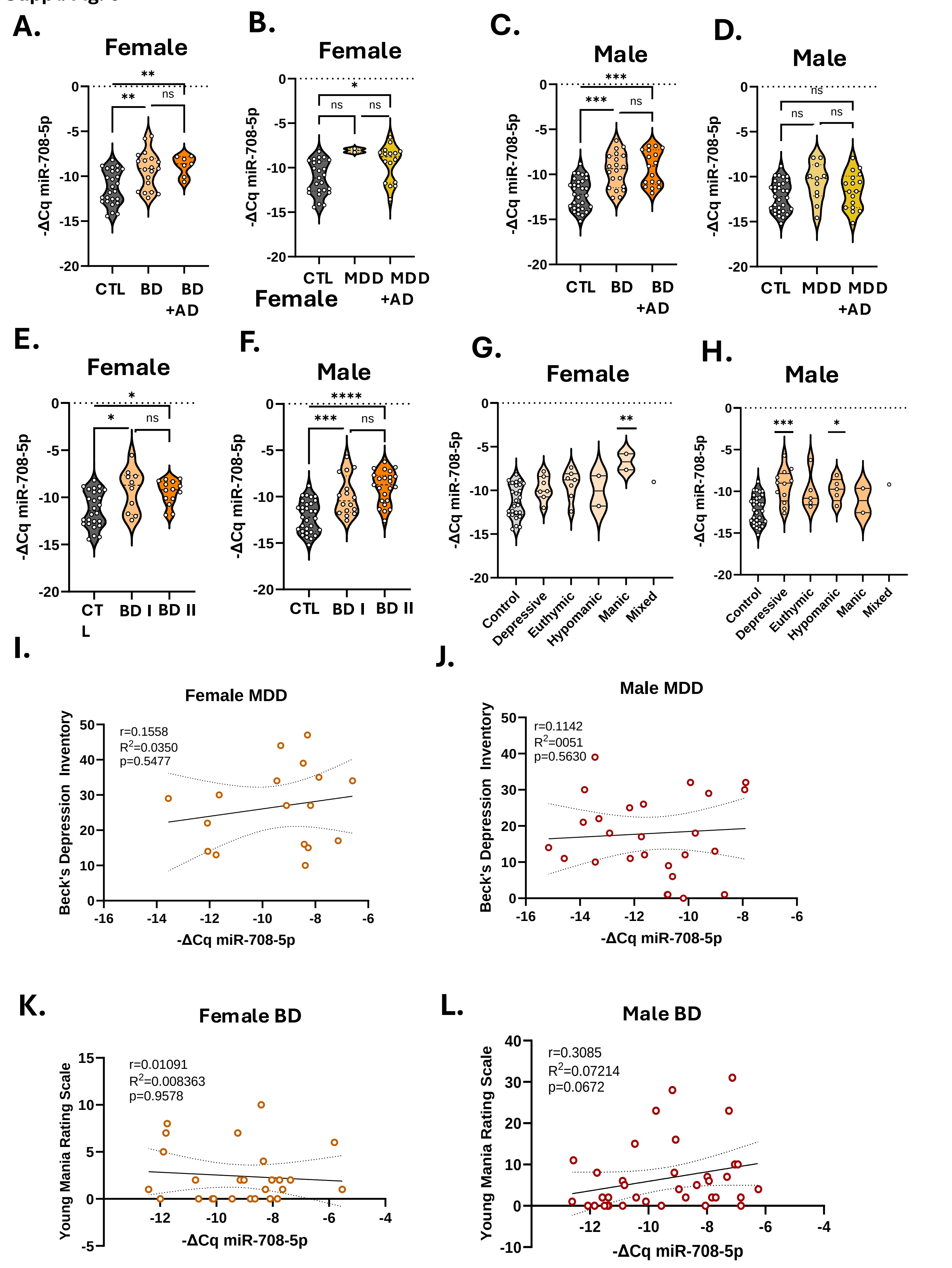
A.
Female
-ΔCq miR-708-5p
CTL
BD
BD +AD
B.
Female
CTL
MDD
MDD +AD
C.
Male
D.
Male
E.
Female
CTL
BD I
BD II
F.
Male
G.
Female
Control
Depressive
Euthymic
Hypomanic
Manic
Mixed
H.
Male
I.
Female MDD
r=0.1558
R2=0.0350
p=0.5477
Beck's Depression Inventory
J.
Male MDD
r=0.1142
R2=0051
p=0.5630
K.
Female BD
r=0.01091
R2=0.008363
p=0.9578
Young Mania Rating Scale
L.
Male BD
r=0.3085
R2=0.07214
p=0.0672

**Suppl. Fig. 7**

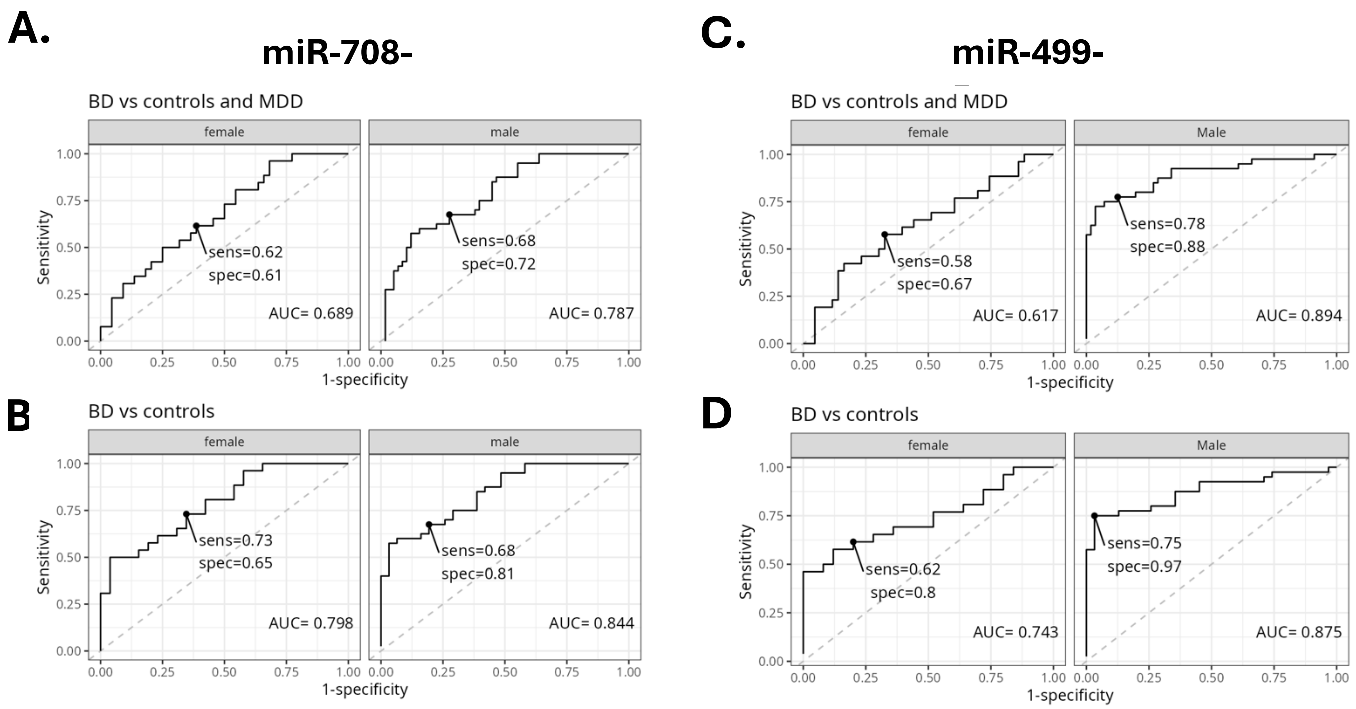

A.
miR-708-
BD vs controls and MDD
female
male
Sensitivity
sens=0.62
spec=0.61
AUC= 0.689
sens=0.68
spec=0.72
AUC= 0.787
1-specificity
B
BD vs controls
female
male
sens=0.73
spec=0.65
AUC= 0.798
sens=0.68
spec=0.81
AUC= 0.844
C.
miR-499-
BD vs controls and MDD
female
Male
sens=0.58
spec=0.67
AUC= 0.617
sens=0.78
spec=0.88
AUC= 0.894
D
BD vs controls
female
Male
sens=0.62
spec=0.8
AUC= 0.743
sens=0.75
spec=0.97
AUC= 0.875
